# Comparing different computational approaches for detecting long-term vertical transmission in host-associated microbiota

**DOI:** 10.1101/2022.08.29.505647

**Authors:** Benoît Perez-Lamarque, Hélène Morlon

## Abstract

Long-term vertical transmissions of gut bacteria are thought to be frequent and functionally important in mammals. Several phylogenetic-based approaches have been proposed to detect, among species-rich microbiota, the bacteria that have been vertically transmitted during a host clade radiation. Applied to mammal microbiota, these methods have sometimes led to conflicting results; in addition, how they cope with the slow evolution of markers typically used to characterize bacterial microbiota remains unclear. Here, we use simulations to test the statistical performances of two widely-used global-fit approaches (ParaFit and PACo) and two event-based approaches (ALE and HOME). We find that these approaches have different strengths and weaknesses depending on the amount of variation in the bacterial DNA sequences and are therefore complementary. In particular, we show that ALE performs better when there is a lot of variation in the bacterial DNA sequences, whereas HOME performs better when there is not. Global-fit approaches (ParaFit and PACo) have higher type-I error rates (false positives) but have the advantage to be very fast to run. We apply these methods to the gut microbiota of primates and our results suggest that only a small fraction of their gut bacteria is vertically transmitted.

## Introduction

Most mammals strongly rely on their associated microbial communities, called microbiota, for various functions like their nutrition, protection, or development (Hacquard et al., 2015; McFall-Ngai et al., 2013; Selosse, Baudoin, & Vandenkoornhuyse, 2004). A range of strategies have evolved to ensure the efficient transmission of some microbes across each generation, including direct transmissions at birth, during parental care, or through social contact (Moran, Ochman, & Hammer, 2019). If these transmissions are stable and faithful, host-microbe interactions are conserved in the host lineage over long-time scales, and we refer to this process as vertical transmission (following the definition of Groussin et al., 2017). At host speciation, vertically transmitted microbes can be inherited by the two daughter host species and separately evolve as independent strains in each host lineage, resulting in a pattern of cophylogeny, where the tree of microbial strains mirrors the host phylogenetic tree (de Vienne et al., 2013; Page, 1994). Conversely, if a microbe is acquired from the environmental pool of microbes at each host generation and if these microbial pools are not as geographically structured as the host species, we do not expect cophylogenetic patterns between the microbe and the host. Cophylogenetic patterns of vertically transmitted microbes can also be erased by frequent horizontal transfers from particular host lineages to others (*i*.*e*. host-switches).

Several studies have reported long-term vertical transmissions among the bacterial gut microbiota of mammals (Gaulke et al., 2018; Groussin et al., 2017; Moeller et al., 2016; Perez-Lamarque & Morlon, 2019; Sanders et al., 2014; Youngblut et al., 2019). Evidence mainly comes from analyses of DNA metabarcoding datasets, where the whole bacterial communities are characterized using the 16S rRNA gene, a short and slowly evolving region (Ochman et al., 2010). A general approach to identifying vertically transmitted bacteria consists in (i) clustering the 16S rRNA sequences into operational taxonomic units (OTUs) based on sequence similarity, (ii) reconstructing for each bacterial OTU a tree of its strains (*i*.*e*. distinct haplotype sequences), and (iii) inferring which OTUs present a cophylogenetic pattern with the host, which would suggest that the corresponding OTUs are vertically transmitted. This general approach has led to estimates of the proportion of vertically transmitted gut bacteria in mammals ranging from more than 50% of all bacterial OTUs (Groussin et al., 2017) to only 14% (Gaulke et al., 2018), or even as few as ∼8% in great apes (Perez-Lamarque & Morlon, 2019). These discrepancies likely come from the use of different approaches to identify cophylogenetic patterns, and from differences in the statistical performances of these different approaches.

Various approaches have been developed in the past decades to identify cophylogenetic patterns, *i*.*e*. to assess the congruence between host and symbiont phylogenies (de Vienne et al., 2013; Dismukes, Braga, Hembry, Heath, & Landis, 2022; Legendre, Desdevises, & Bazin, 2002; Page, 1994). These approaches have been applied to a variety of host-symbiont systems and have advanced our understanding of the evolution of such symbiotic interactions (Blasco-Costa, Hayward, Poulin, & Balbuena, 2021; Hayward, Poulin, & Nakagawa, 2021). In particular, cophylogenetic approaches have been used to assess whether and how host-symbiont associations impact deep-time evolutionary processes, through vertical transmission and/or preferential host-switches (Blasco-Costa et al., 2021; De Vienne, Giraud, & Shykoff, 2007; de Vienne et al., 2013). A common difficulty of these interpretations is the fact that shared biogeographic structure between hosts and symbionts, for example as a result of vicariance, can also generate cophylogenetic patterns. In particular, if environmental pools of symbionts differ across geographic regions occupied by closely related host species, this can generate cophylogenetic patterns in the absence of vertical transmission (Amato et al., 2019; Perez-Lamarque, Krehenwinkel, Gillespie, & Morlon, 2022). Additional precautions are therefore required to link a cophylogenetic pattern to long-term vertical transmission.

Co-phylogenetic approaches can be divided into two main categories (de Vienne *et al*., 2013, Table 1). The first category, referred to as ‘global-fit’ approaches, measures a global congruence between the host and symbiont evolutionary histories. For instance, ParaFit (Legendre et al., 2002) and PACo (Balbuena, Míguez-Lozano, & Blasco-Costa, 2013) are two widely used approaches based on the fourth-corner statistic or Procrustes superimposition, respectively. These approaches can be directly applied to the symbiont genetic distances and thus do not require a robust reconstruction of the symbiont phylogenetic tree. They can also handle multiple strains per extant host species. However, they only provide a measure of a cophylogenetic pattern and do not inform on the processes at play. The second category of approaches, referred to as ‘event-based’ approaches, directly models the events of codivergence, host-switches, duplications, and/or losses, to reconciliate the host and symbiont phylogenies, while considering the uncertainly in the symbiont evolutionary history (Figure 1). For instance, ALE (Szöllősi, Rosikiewicz, et al., 2013) uses a posterior distribution of symbiont phylogenetic trees to fit reconciliation events. This approach therefore fully models phylogenetic uncertainty in contrast with mainstream event-based approaches using maximum parsimony, e.g. TreeMap (Page, 1994) or eMPRess (Santichaivekin et al., 2021), which are better suited when phylogenetic reconstructions are robust. ALE also accounts for the possibility that the symbiont was absent in the ancestor of all hosts and only secondarily acquired (Szöllősi, Tannier, et al., 2013). It considers unsampled or extinct host lineages and even assumes that host-switching between sampled lineages always involves an unsampled or extinct host lineage as an intermediate (Szöllősi, Tannier, et al., 2013). This is particularly relevant as under-sampling of the extant host species is often important when looking at dynamics of microbial transmission in large animal clades such as mammals (Groussin et al., 2017; Youngblut et al., 2019)). A limitation of ALE is that only uses the information contained in the topology of phylogenetic trees, and sometimes the order of the nodes in the host phylogenetic tree, but not branch lengths (Szöllősi, Tannier, et al., 2013). Another approach (called HOME) was recently developed with the specific aim of analyzing microbiota (meta)barcoding data with little phylogenetic information by avoiding the reconstruction of unreliable trees of OTU strains (Perez-Lamarque & Morlon, 2019). Rather than first reconstructing trees of OTU strains as in ALE, HOME directly models the bacterial DNA substitution process on the host phylogeny under a scenario of vertical transmission with potential host-switches. A limitation of the approach is that it cannot handle multiple OTU strains per extant host, as it does not model duplication events and assumes replacement of the microbial strain rather than coexistence upon host-switch. HOME does not explicitly model potential losses of microbial strains either, nor does it consider unsampled or extinct host lineages. Yet, the direct modeling of DNA evolution offered by HOME can be particularly valuable when the host clade has diverged recently and molecular markers used to characterize the microbiota have accumulated very few substitutions.

**Table 1.** Main characteristics of the different cophylogenetic methods that can be used for detecting vertical transmission in host-associated microbiota. ParaFit, PACo, and ALE were developed to study a broader array of host-symbiont associations. HOME was specifically designed to study host-microbiota associations from short (meta)barcoding data.

| Method | ParaFit | PACo | ALE | HOME |
| --- | --- | --- | --- | --- |
| **Main feature** | **Global-fit approach:**<br>Measures the overall congruence between the host phylogeny and the microbial genetic distances using the fourth-corner statistic. | **Global-fit approach:**<br>Measures the overall congruence between the host phylogeny and the microbial genetic distances using Procrustes superimposition. | **Event-based approach:**<br>Models events of codivergence, host-switch, loss, and duplication to reconcile the host and microbial phylogenetic trees (reconstructed beforehand). | **Event-based approach:**<br>Models the microbial DNA evolution on the host phylogenetic tree, while fitting potential host-switches. |
| **Major assumptions** | Identical OTU strains in different host species are considered as interchangeable. | Identical OTU strains in different host species are considered as interchangeable. | Assumes that most host lineages are not sampled or extinct;<br>A host-switch always involves an unsampled lineage as an intermediate.<br>Branch lengths are not considered (only the topologies and the relative order of nodes of the host tree are considered). | Models DNA evolution on the host phylogenetic tree, allowing for host-switches.<br>OTU duplications and losses are not considered. Does not consider under-sampling or extinction of the host lineages. |
| **Inputs** | Pairwise (phylo)genetic distances for the hosts and their associated microbes | Pairwise (phylo)genetic distances for the hosts and their associated microbes | A rooted host phylogenetic tree and a posterior distribution of microbial phylogenetic trees (with at least one microbe per host species) | A time-calibrated host phylogenetic tree and a DNA alignment constituted of one microbial DNA strain per host species (at most) |
| **Randomization strategy** | **Null model 1:** Independent permutation of the host species associated with each microbial strain<br>**Null model 2:** Permutation of the host species name in the host phylogenetic tree | **Null model 1:** Independent permutation of the host species associated with each microbial strain<br>**Null model 2:** Permutation of the host species name in the host phylogenetic tree | **Null model 2:** Permutation of the host species name in the host phylogenetic tree | **Null model 2:** Permutation of the host species name in the host phylogenetic tree |

**Figure 1:**
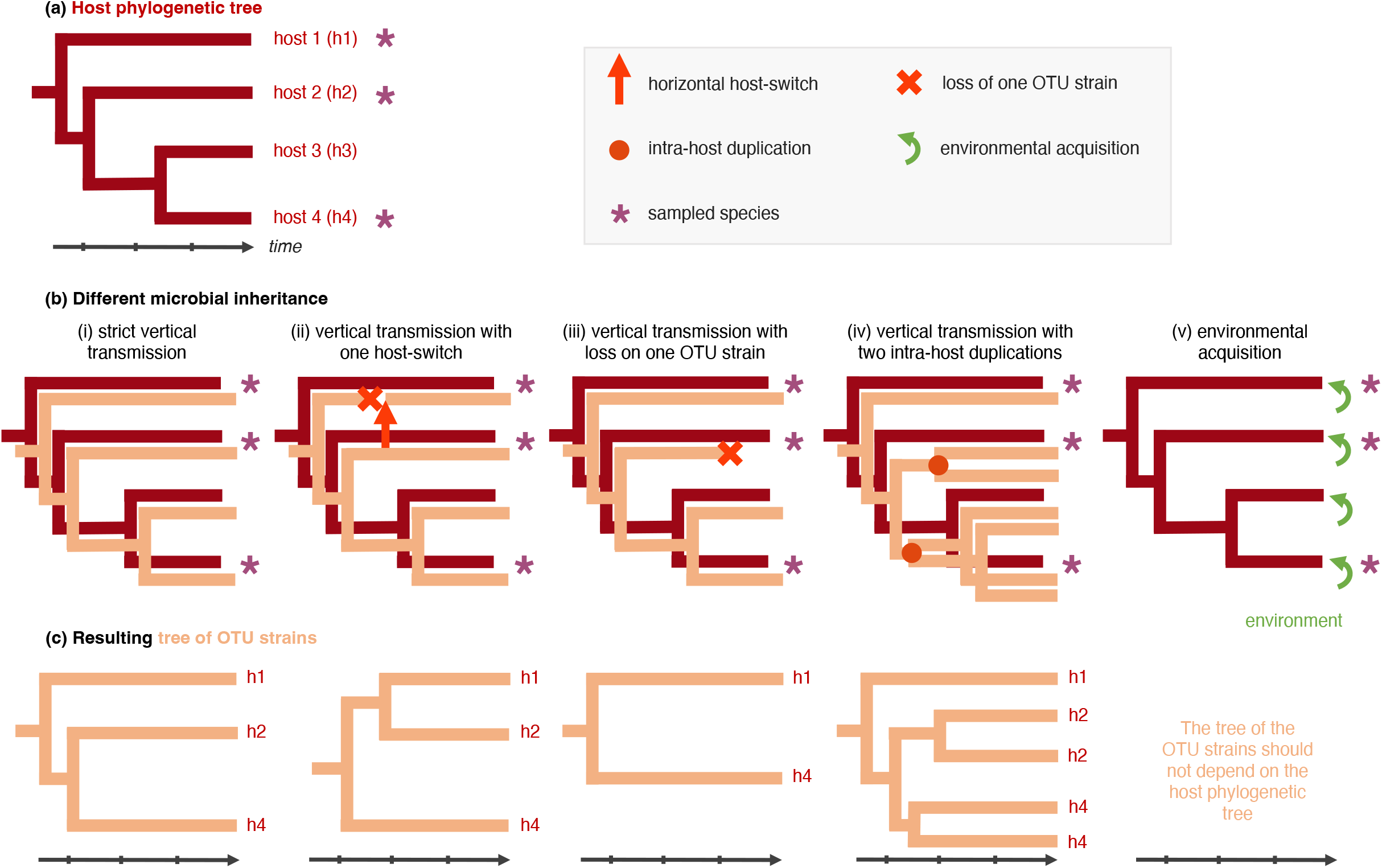
Different modes of inheritance of a given host-associated operational taxonomic unit (OTU) and their consequences on the microbial tree (tree of OTU strains). On a phylogenetic tree of 4 host species (a), we represent the different modes of inheritance for a given OTU (b) and the resulting tree of OTU strains (c). Each host is at least colonized by one strain of this given OTU (except in the case of OTU loss in (iii)). We also represent a sampling process where the microbiota of only some extant host species (marked by “*”) is characterized. Extreme scenarios correspond to strict vertical transmission (i; perfect cophylogenetic pattern) or environmental acquisition (v; no cophylogenetic pattern expected). In more intermediate scenarios, the perfect congruence between the host phylogeny and the tree of OTU strains, a characteristic of vertical transmission, is dampened by events of horizontal transmissions (ii; the horizontal host-switch from one donor host to a receiver host with replacement of the OTU strain), microbial loss (iii), or intra-host duplication (iv).

Here, we aim to test the statistical performances of different cophylogenetic approaches to detect vertical transmission in microbiota characterized by metabarcoding techniques. We simulate the evolution of 16S rRNA gene sequences for bacteria that are either vertically transmitted or evolving independently of the host phylogeny. We measure the statistical power (the proportion of vertically transmitted bacteria inferred as being vertically transmitted) and the type-I error rate (the proportion of independently evolving bacteria inferred as being vertically transmitted; *i*.*e*. false-positives) of ParaFit, PACO, ALE, and HOME. Then, we apply these different methods to the gut bacterial microbiota of 18 new-world and old-world primate species using the dataset generated by Amato et al. (2019). Based on our results, we discuss the pros and cons of each approach and highlight promising areas for future development.

## Methods

### Primate phylogeny

In order to design our simulations with realistic settings and to better interpret our empirical analyses of the gut microbiota of primates (Amato *et al*. (2019)), we performed all the simulations on the primate phylogenetic tree of Dos Reis *et al*. (2018). This tree is a nearly complete phylogenetic tree of extant primates (367 species) reconstructed using phylogenomic data and fossil calibrations, with a crown age estimate of ∼74 million years (Myr). For computational reasons, we scaled the age of the primate phylogeny to 1 (relative timing) using the R-package ape (Paradis, Claude, & Strimmer, 2004; R Core Team, 2022).

### Simulations

We simulated three different scenarios of host-microbiota evolution on the complete primate phylogenetic tree (Figure 1): (I) strict vertical transmission where each microbial OTU evolves directly on the host phylogeny (Figure 1b i), (II) vertical transmission with a given number of horizontal host-switches (5, 10, 15, or 20, Figure 1b ii), or (III) environmental acquisition, where the microbes evolve independently and are randomly acquired by the extant host species (Figure 1b v). Each simulation generates a tree of OTU strains (Figure 1c). For II (vertical transmission with host-switches), we considered that host-switches can happen uniformly on the host phylogeny from a donor branch to a receiving branch where it replaces the previous OTU strain (Figure 1b ii). The range of simulated horizontal host-switches was chosen to test their effect when they were rare to moderately frequent, as codivergence with very frequent switches can no longer be considered as a scenario of vertical transmission. For III (environmental acquisition), the tree of an independently-evolving OTU was obtained by simulating a pure birth process using the function *pbtree* (R-package ape (Paradis et al., 2004)) until reaching as many tips as primate species; we then randomly assigned each tip in the tree of OTU strains to a primate species, mimicking the process of random strain acquisition from an environmental pool (Figure 1b v).

For each scenario and each tree of OTU strains, we simulated on this tree the evolution of a 300 bp DNA region mimicking the V4 region of the 16S rRNA gene. We thus obtained for each OTU a DNA alignment made of the DNA sequences from each extant host species. We assumed that 10% of the sites were variable (other sites are kept conserved) and for these variable sites, DNA substitutions were modeled using a K80 process (Kimura, 1980) with different relative substitution rates (μ): 1.5 (many substitutions), 1, 0.5, 0.1, and 0.05 (very few substitutions). These relative rates were chosen to obtain numbers of segregating sites and strains in the simulated alignments that are consistent with the empirical within-OTU alignments: For μ=1.5, we obtained alignments with a mean number of segregating sites >20 and a total number of strains >15, while for μ=0.05, the simulated alignments had on average <5 segregating sites and <5 number of strains (Supplementary Fig. 1). These simulations were performed using the function *sim_microbiota* in the R-package HOME (Perez-Lamarque & Morlon, 2019; R Core Team, 2022).

Finally, to insert our simulations in the frequently encountered situation when only a small fraction of the extant host species has their microbiota characterized, we retained only the simulated OTU strains present in the 18 primate species sampled in Amato *et al*. (2019). For each OTU, a single OTU strain is associated with each host species. We refer to the corresponding alignments as the ‘simulations without duplications or losses’.

In addition to the simulations without duplications or losses, we considered that OTU strains can be lost during host evolution or not detected in extant host-associated microbiota using metabarcoding techniques (Figure 1b iii). To mimick this, we randomly sampled, in each alignment, the strains of 10 out of 18 extant host species. We refer to these alignments as the “simulations with losses”. We also considered that intra-host duplications can happen stochastically during host evolution (Figure 1b iv), such that multiple OTU strains can persist in a host lineage. We simulated the same scenarios as above, but simultaneously simulated duplication events using a continuous-time Markov process, *i*.*e*. duplications can happen at any time on the host branches, with a relative rate δ=2. We obtained alignments by selecting the OTU strains present in each of the 18 primate species. We referred to these alignments as the “simulations with duplications.” Finally, we simulated losses and/or non-detection in the simulations with duplications, by randomly sampling the OTU strains of 10 out of 18 extant host species in each alignment. We thus obtained “simulations with losses and duplications”.

For each simulated scenario and combination of parameters, we performed 100 simulations. We therefore obtained a total of 12,000 simulated alignments.

### Inferring vertically transmitted OTUs

We considered four different approaches for detecting vertical transmission: two global-fit approaches, ParaFit and PACo, and two event-based approaches, ALE and HOME (Table 1). Other global-fit approaches exist for detecting vertical transmission, like the global-fit approaches proposed by Hommola *et al*. (2009), which is a generalization of the Mantel tests, but we chose to only focus on the two most frequently used ones (ParaFit and PACo: Gaulke et al., 2018; Youngblut et al., 2019).

Event-based and global-fit approaches rely on the same randomization-based approach to assess statistical significance. After fitting the model under a scenario of vertical transmission (for event-based approaches) or computing the statistic of the test (for global-fit approaches), randomizations are used for generating null expectations under a scenario of independent host-OTU evolution. By comparing the observed fit to null expectations, we may reject the null hypothesis of independent evolution and conclude that the OTU is vertically transmitted. Two different randomization schemes have been used (Table 1). For ParaFit and PACo, Balbuena et al. (2013) and Legendre et al. (2002) used a randomization scheme that we refer to as ***null model 1*** (also referred to as “r0” in Hutchinson *et al*. (2017)), which consists in permuting the host species associated to each OTU strain, independently for each strain. The number of host species per OTU strain is therefore maintained, but the number of OTU strains per host species is not, and can even reach 0. For ALE and HOME, a stricter randomization scheme has been used (that we refer to as ***null model 2***) that consists in shuffling species names in the host phylogenetic tree, which guarantees that each host species has at least one OTU strain. We used this null model in our ALE and HOME analyses and used it also in addition to ***null model 1*** for ParaFit and PACo (Table 1). The choice of the number of randomizations used for these tests results from a trade-off between accuracy and computation time. Given the computational requirements of the different approaches (see Results), we used 10,000 randomizations for the global fit approaches (ParaFit and PACo) and 100 for the event-based approaches (ALE and HOME).

We ran ParaFit and PACo on the phylogenetic distances between pairs of extant primate species and the microbial genetic distances between pairs of OTU strains. We computed these genetic distances using a K80 model of DNA substitution, which corrects for potential mutation saturation. ParaFit and PACo statistics were both computed using a Cailliez correction for negative eigenvalues. We amended the functions *parafit* and *PACo* from the R-packages ape (Paradis et al., 2004) and paco (Hutchinson, Cagua, Balbuena, Stouffer, & Poisot, 2017) respectively, to avoid technical issues when the number of OTU strains is low. To evaluate the significance of the statistic of each test of vertical transmission, we compared its value to a null distribution under the hypothesis of independent host-OTU evolution using 10,000 randomizations, with both ***null model 1*** and ***null model 2***. Balbuena et al. (2013) recommend using 100,000 permutations for high precision; we reduced this number here to save computational time and energy and checked that this did not affect our results on a subset of simulations (see Results). To avoid computational issues during the randomizations of the associations between hosts and OTU strains, we only ran ParaFit and PACo for the alignments containing at least 3 different strains.

To run ALE, one needs first to generate a posterior distribution of trees of OTU strains using Bayesian phylogenetic inference. We reconstructed phylogenetic trees for each alignment using PhyloBayes (Lartillot & Philippe, 2004) following Groussin *et al*. (2017). We ran PhyloBayes for 4,000 generations, sampling at every generation after an initial burn-in of 1,000 generations. We then ran ALE with the host phylogeny and the distribution of trees of OTU strains as inputs, using the *ALEml* program available at https://github.com/ssolo/ALE. ALE estimates the maximum likelihood rates of host-switch, duplication, and loss, and generates a set of host-OTU reconciliations. We used 100 reconciliations and computed an average number of codivergences, host-switches, duplications, and losses. To evaluate the significance of these estimated scenarios of vertical transmission, we shuffled the primate species in the phylogenetic tree (***null model 2***) and re-ran ALE to obtain a distribution of the number of reconciliation events under a null hypothesis of independent host-OTU evolution. We then compared two criteria to reject the null hypothesis. First, we used the criterium of Groussin *et al*. (2017): (i) the estimated number of codivergences is significantly higher than the number of host-switches and (ii) under a null hypothesis of independent host-OTU evolution, the estimated number of codivergences is higher than the number of host-switches in at most 5% of the null expectations. Second, as in Dorrell *et al*. (2021), we considered that an OTU is vertically transmitted if (i) the estimated number of codivergences is higher than 95% of the null expectations and if (ii) the estimated number of host-switches is lower than 95% of the null expectations. We performed 100 randomizations per OTU, except when analyzing simulations with intra-host duplications; in this case, ALE is more computationally intensive and we thus performed only 50 randomizations. We ran ALE only for alignments that had at least one segregating site.

We ran HOME using the function *HOME_model* in the R-package HOME (Perez-Lamarque & Morlon, 2019). For each alignment, HOME outputs the maximum-likelihood estimates of the number of host-switches and the substitution rate. In HOME, the likelihood is estimated using Monte Carlo simulations (Perez-Lamarque & Morlon); here we used 5,000 simulated trees and picked the tested numbers of host-switches in a grid from 1 to 35. Because we simulated the process of host-switching on the complete primate phylogeny (367 species) and that HOME can only estimate host-switches occurring between lineages present in the reconstructed phylogenetic tree (composed of only 18 species), we expected HOME to infer fewer switches than simulated (Table 1). As for ALE, we assessed the significance of the estimated scenario of vertical transmission by performing 100 randomizations shuffling the associations between host and OTU strains (***null model 2***). We considered that an OTU was vertically transmitted if both the estimated substitution rate and the observed number of host-switches were lower than 95% of the null expectations. Because HOME does not tolerate multiple OTU strains per host tip at present, when simulations included duplications, we randomly picked one single strain per host species. We ran HOME only for alignments that had at least one segregating site.

We computed the statistical power as the percentage of OTUs simulated as vertically transmitted (strictly vertically transmitted or vertically transmitted with host-switches) that were correctly inferred as being vertically transmitted and the type-I error rate as the percentage of OTUs simulated as independently evolving that were incorrectly inferred as being vertically transmitted. We also measured the computation time of the different approaches using a random subset of simulations. For event-based approaches, we also evaluated the accuracy of parameter estimation.

### Empirical application

We downloaded the dataset from Amato *et al*. (2019) characterizing the gut bacterial microbiota of 153 primates belonging to 18 species using the V4 region of the 16S rRNA gene available at https://www.ebi.ac.uk/ena/data/view/PRJEB22679. The demultiplexed Illumina reads were processed using a pipeline based on VSEARCH (Rognes, Flouri, Nichols, Quince, & Mahé, 2016) available at https://github.com/BPerezLamarque/Scripts/. In short, after quality filtering, the reads were clustered into OTUs using either Swarm clustering (Mahé, Rognes, Quince, de Vargas, & Dunthorn, 2015) or traditional OTU clustering methods with 95% or 97% sequence similarity thresholds using VSEARCH. We tested different OTU clustering methods because we cannot know *a priori* which clustering method will delineate a given OTU at the “right” level (*i*.*e*. not merging two biological units within the same OTU, nor over-splitting one biological unit into two OTUs; Perez-Lamarque & Morlon, 2019). Chimeras were filtered out *de novo* and taxonomy was assigned to each OTU using the Silva database (Quast et al., 2013). We kept only non-chimeric bacterial OTUs represented by at least 5 reads in at least 2 samples. Finally, we assumed that if an OTU had less than 5 reads in a sample, it was likely cross-contamination and set its abundance to 0. For Swarm clustering, we obtained 6,373 OTUs representing a total of 4,019,271 reads, while clusterings at 97% and 95% gave 5,624 and 4,663 OTUs respectively (corresponding to a total of 4,088,586 and 4,300,861 reads).

The ability to detect vertical transmission for a particular OTU depends on the ability to detect this OTU across species in the first place, which can depend on a number of factors during DNA extraction, PCR amplification, and sequencing. We therefore evaluated whether we successfully detected most of the OTUs present within samples and primate species by performing rarefaction analyses using the *vegan* R-package (Oksanen et al., 2016).

We tested the support for vertical transmission only for “core OTUs”, taken to be OTUs present in at least 10 out of the 18 primate species represented in the dataset. First, we built a dataset with only one OTU strain per host species: for each OTU, we merged all the primate samples from the same species together and built the alignment by picking per host species the most abundant strain assigned to this OTU. We aligned OTU strains using MAFFT (Katoh & Standley, 2013). We recorded the number of segregating sites and unique strains in the resulting alignments and we applied ParaFit, PACo, ALE, and HOME to detect vertically transmitted OTUs. Given that our simulations highlighted a high type-I error rate for the correlative approaches and ALE, and a low statistical power for HOME, when the number of segregating sites and the number of hosts are low (see Results), we compared the distribution of these characteristics in OTUs inferred to be vertically transmitted or not by the different approaches. A comparatively low number of segregating sites and/or hosts in OTUs inferred to be vertically transmitted by the correlative approaches and ALE would suggest a lot of false positives. A comparatively high number of segregating sites and/or hosts in OTUs inferred to be vertically transmitted by HOME would suggest that some vertically transmitted OTUs might be missed (false negatives).

Next, we relaxed the hypothesis of a single OTU strain per host species, by considering the possibility of multiple strains, resulting for instance, from intra-host duplications: we picked up to 3 OTU strains per host species by selecting the 3 most abundant ones when available. Given that HOME cannot tolerate multiple OTU strains per host, we only ran ParaFit, PACo, and ALE.

Finally, for the OTUs that presented a significant cophylogenetic pattern according to the different approaches, we tested whether this cophylogenetic pattern could come from a geographic effect rather than from vertical transmission. Indeed, as noted by Amato *et al*. (2019), a cophylogenetic pattern in primate microbiota could arise because of the geographic split of the primates between the Old World (Africa and Asia) and the New World (Americas). Heterogeneous environmental pools of microbes in the Old and New Worlds combined with the fact that closely related primate species tend to be present in the same area could generate cophylogenetic patterns in the absence of vertical transmission. To test this, we randomized the associations between primate species and OTU strains within the Old World and New World respectively, and re-ran the analyses. If we still detect a significant cophylogenetic pattern, we cannot reject the hypothesis that this pattern (at least partially) comes from heterogeneous pools of microbes between the Old World and the New World. Conversely, if we no longer detect a significant cophylogenetic pattern, we can reject this hypothesis, therefore suggesting that the cophylogenetic pattern is linked to vertical transmissions. An alternative (and faster) way for event-based approaches to test this would be to use post-processing of the inferences to assess whether the host-switches inferred by ALE and HOME tend to be more frequent between host lineages present on the same continents (Perez-Lamarque et al., 2022). Here, for the sake of comparisons between global-fit and event-based approaches, we evaluated the hypothesis of different geographic pools of microbes using randomizations.

## Results

### Computational efficiency

Global-fit approaches, especially ParaFit, were the fastest (Table 2). Their computation time using 10,000 randomizations increased slightly with higher simulated substitution rates (μ), but remained on average lower than one minute (when measured on an Intel 2.8 GHz MacOSX laptop using only 1 CPU; Table 2; Supplementary Fig. 2). In contrast, both event-based approaches were much slower, even though we used only 100 randomizations to evaluate their significance. The computation time of HOME increased with increasing μ; from only a few hours when the number of segregating sites was very low, to several hours or a few days when there were many of them (Table 2). The computation time of ALE increased with decreasing μ, linked to an increase of phylogenetic uncertainty in the trees of OTU strains that slows down the reconciliation between these trees and the host phylogeny. ALE sometimes took several days to run for a single OTU with μ=0.05 (Table 2; Supplementary Fig. 2). It also significantly increased in the presence of duplications.

**Table 2.**
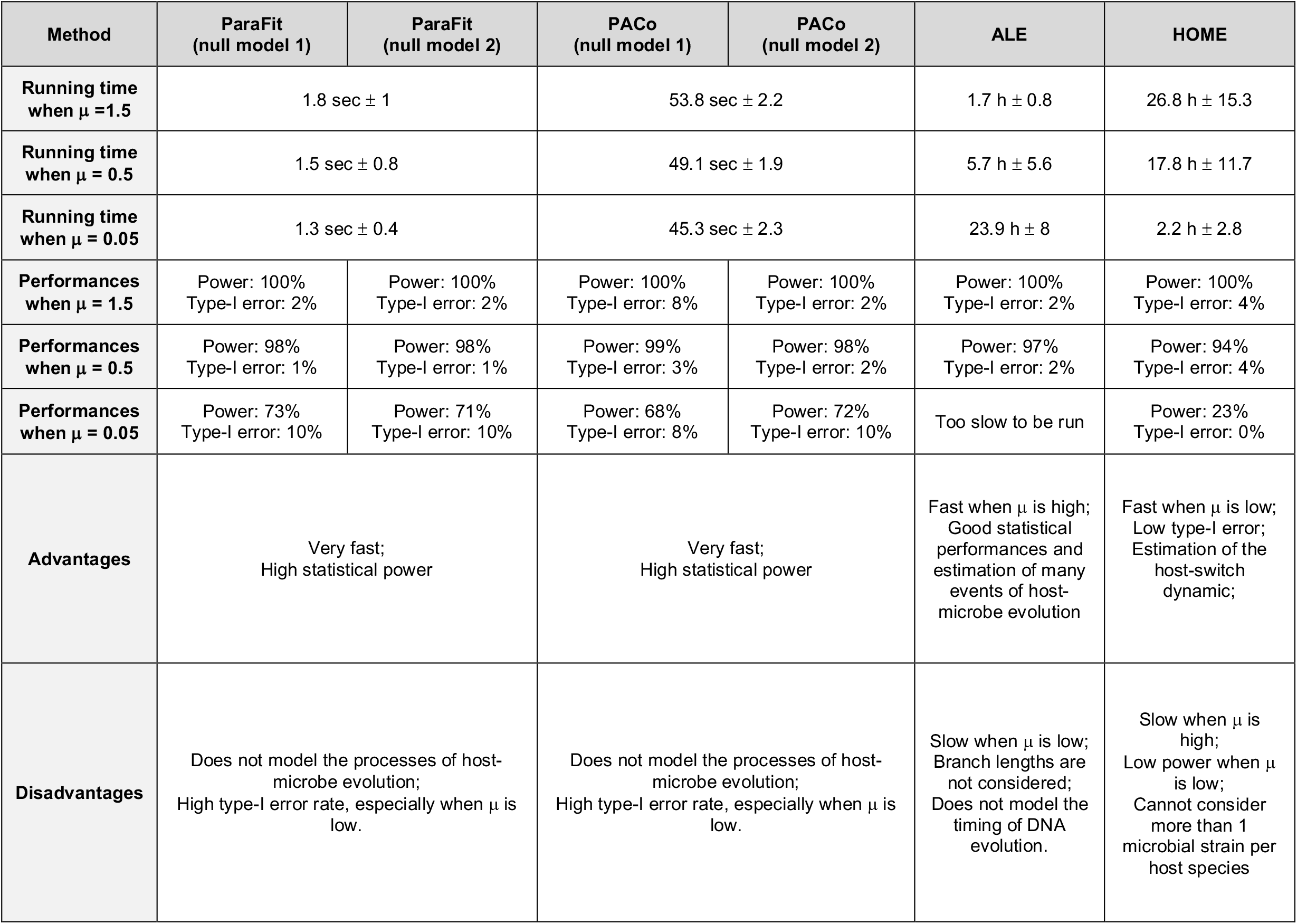
Summary of the statistical performances of the different methods for detecting vertical transmission in host-associated microbiota: For each method, we summarize its running time and its statistical performances evaluated using simulations. The running time and statistical performances correspond to the simulations without duplications or losses, with different substitution rates for the microbial OTUs: μ=1.5, μ=0.5, and μ=0.05. Global-fit approaches were run with 10,000 randomizations, while we only used 100 randomizations for event-based ones. Computation times (mean ± s.d.) were measured on an Intel 2.8 GHz MacOSX laptop using only 1 CPU; for ALE, they included the time to reconstruct the trees of OTU strains (e.g. using PhyloBayes), which is longer when the substitution rate is high. The inference of HOME was designed to run in parallel and is thus faster on multi-core processors. Details on the statistical performances of the different approaches can be found in the supplements.

**Figure 2:**
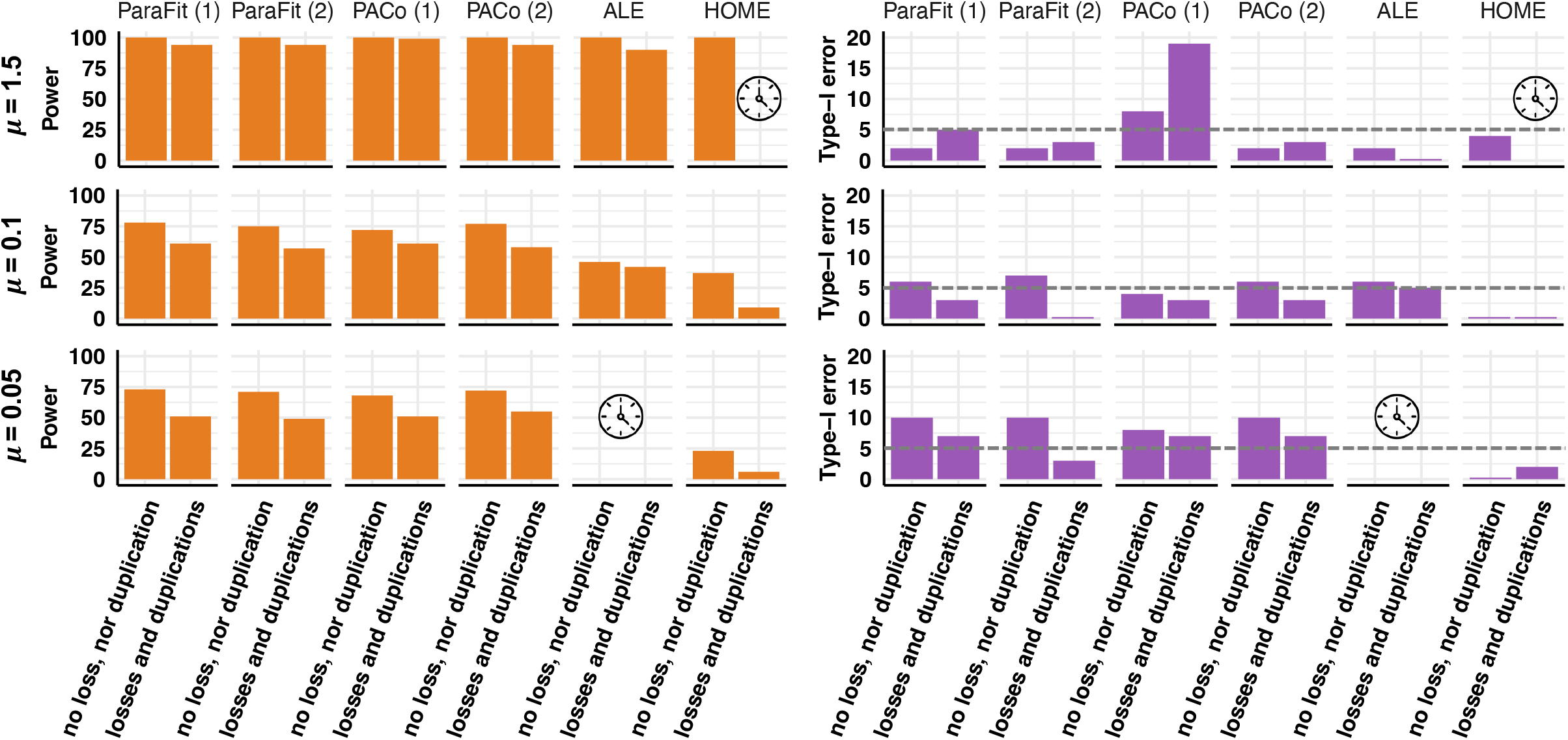
Summary of the statistical performances of the different methods for detecting vertical transmission in host-associated microbiota: For each method, we indicated the statistical power (left) and the type-I error rate (right) for different substitution rates for the simulated microbial OTUs: μ=1.5, μ=0.1, and μ=0.05. Statistical performances are reported for simulations without duplications or losses and simulations with both duplications and losses. The clock symbol indicates analyses that were too computationally intensive to be run in a reasonable amount of time and energy expense. Horizontal dashed grey lines indicate a 5% type-I error rate. Details on the statistical performances of the different approaches can be found in the supplements.

To save time and energy, for each combination of simulated parameters, we ran ALE and HOME on only 50 simulated alignments (against 100 for global-fit approaches), except for μ=0.05, where we used 100, given that many of the resulting alignments contained no segregating sites. In addition, we did not run ALE when μ=0.05 and did not use HOME for alignments simulated with both μ>0.5 and duplications.

### Simulations without duplications or losses

The alignments simulated without duplications or losses contained a mean number of segregating sites larger than 20 and a mean number of strains larger than 15 (almost one OTU strain for each host species) when the simulated substitution rate (μ) equaled 1.5. With μ=0.05, they contained less than 5 segregating sites (with many alignments presenting no segregating sites; Supplementary Fig. 1) and less than 5 strains.

Simulating up to 20 host-switches on the complete primate phylogeny had a limited impact on the statistical performances of the different approaches (see Supplementary Figs. 2-17). Therefore, we hereafter pooled all simulations of vertical transmission (*i*.*e*. strict or with host-switches) when reporting estimations of statistical power.

We found that global-fit approaches (ParaFit and PACo) have a high statistical power (≥98%) when μ≥0.5 regardless of the null model used to assess statistical significance (Figure 2). Their power decreases to ∼70% when μ=0.05 (Supplementary Figs. 3 & 4). However, they also have a rather elevated type-I error rate when μ=0.05 (type-I error rate ∼10% for both ParaFit and PACo). With null model 1, PACo has a type-I error rate >5% even when μ is high (Supplementary Fig. 4).

**Figure 3:**
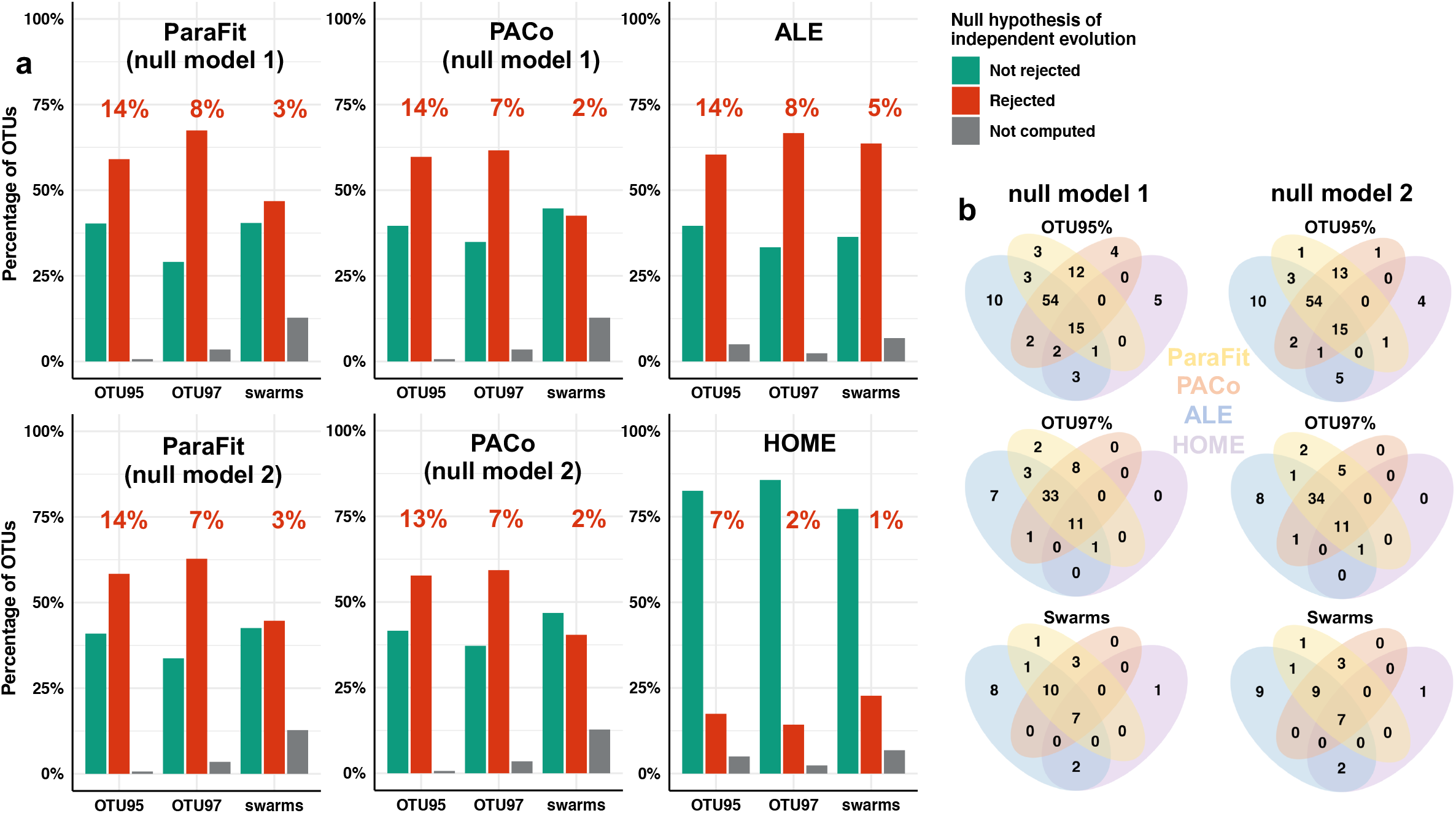
Evidence for vertical transmission in primate gut microbiota varies according to the different methods used. (a) Percentage of core OTUs from the gut microbiota of primates rejecting (in red) or not (in green) the null hypothesis of independent evolution according to the different approaches tested: ParaFit (with null models 1 or 2), PACo (with null models 1 or 2), ALE, or HOME. OTUs colored in red thus represent vertically transmitted OTUs. Percentages at the top of each bar indicate the percentage of reads corresponding to these vertically transmitted OTUs in the whole primate gut microbiota. OTUs were clustered using the 95% or 97% similarity threshold or as Swarm OTUs. (b) Venn diagrams indicating the number of OTUs that are simultaneously inferred to be vertically transmitted using the different approaches.

**Figure 4:**
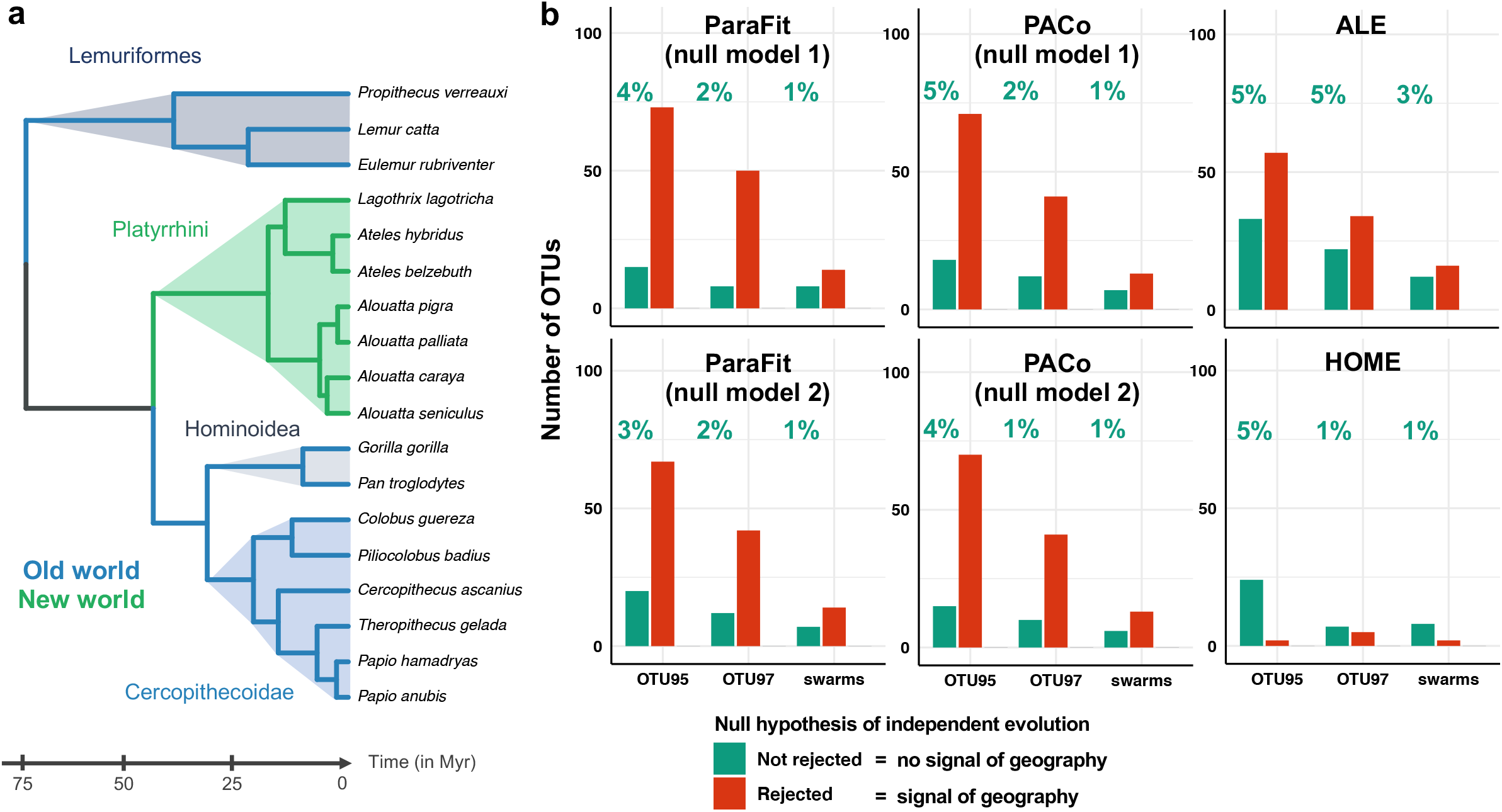
Geography-driven cophylogeny: (a) Phylogenetic tree of the 18 primates with branches colored according to their native geographic area (the New or Old World). (b) Number of OTUs, within those inferred to be vertically transmitted according to ParaFit (with null models 1 or 2), PACo (with null models 1 or 2), ALE, or HOME, that reject (in red) or not (in green) the hypothesis of independent evolution after randomizing the associations between primate species and OTU strains within the Old and New Worlds. If the cophylogenetic pattern is still significant (in red), the original cophylogenetic signal is at least in part driven by geography (i.e. heterogeneous environment pools of bacteria). If the cophylogenetic pattern is no longer significant (in green), the original cophylogenetic signal arises from vertical transmissions (or other, non-geographic effects). At the top of each bar, we indicated the percentage of reads corresponding to these vertically transmitted OTUs in the whole primate gut microbiota.

ALE has a high power (>95%) and a low type-I error (<5%) when μ≥0.5, using either the criteria of Dorrell et al. (2021) or Groussin et al. (2017) for rejecting the null hypothesis of independent evolution (Figure 2; Supplementary Figs. 5a and 6). In simulations with strict vertical transmission, it correctly infers exclusively codivergence events (and their approximate number, *i*.*e*. 17 on an 18 species tree, Supplementary Fig. 5b). ALE also infers host-switches when simulated, although their number is underestimated (Supplementary Fig. 5b). In simulations with independent evolution, *i*.*e*. when there is no ‘correct’ reconciliation scenario, ALE estimates a lower number of codivergence events and a much higher number of host-switches, as expected (Supplementary Fig. 5b). However, with less segregating sites (μ=0.1), the power of ALE is below 50%, the type-I error increases to 6%, and the inference of reconciliation events is not accurate (Supplementary Fig. 5a&b and 6). Indeed, ALE estimates many spurious losses and hosts-switches in simulations with strict vertical transmission, and underestimates the number of host-switches in simulations with host-switches (Supplementary Fig. 5b). The statistical power of ALE is even lower (below 20% for μ=0.1, Supplementary Fig. 6) when following the criterium of Groussin et al. (2017) than that of Dorrell et al. (2021); we thus kept the latter for rejecting the null hypothesis of independent evolution when using ALE in the following analyses.

HOME has a high power (>94%) when μ≥0.5 (Figure 2; Supplementary Fig. 7a), but its statistical power decreases a lot with small μ values: with μ=0.1, the power of HOME is below 40%, and with μ=0.05 below 25%. HOME has a low type-I error rate (<5%) in all conditions, including when μ is low (Supplementary Fig. 7a). In terms of inferred parameters, HOME correctly estimates the substitution rate. We cannot directly test if the number of host-switches is well recovered, as HOME infers switches on the provided tree (with 18 species) rather than on the complete tree, but we find that HOME infers more host-switches when more are simulated (at least with sufficient segregating sites, Supplementary Fig. 7b). In simulations with independent evolution, *i*.*e*. when the evolution of the microbial DNA sequences on the host phylogeny fits poorly, HOME estimates both high substitution rates and a high number of host-switches, as expected (Supplementary Fig. 7b).

### Simulations with losses or/and duplications

When simulating losses (or non-detection within hosts), the statistical power of all the approaches decreases (Supplementary Figs. 8-10), especially for HOME (∼10% when μ=0.05). For simulations with low μ values, the type-I error rate increases strongly (>10%) for global-fit approaches and ALE, but not HOME (0% when μ=0.05). The type-I error rates of global-fit approaches decrease when using null model 2 instead of null model 1 (Supplementary Figs. 8).

Simulations with duplications generated alignments with a higher number of segregating sites and strains (Supplementary Fig. 11). Under this scenario, we found that the statistical power of global-fit approaches remains very high (>60% for all μ; Supplementary Fig. 12). The type-I error rate increases strongly under null model 1, reaching 20% for PACo when μ=1.5, but it can be reduced to 5% by using null model 2. ALE handles duplications very well, conserving a high power (>95%) and a low type-I error rate (<5%; Supplementary Fig. 13). However, the computation time of the approach increases substantially, which complicates the use of the method when the number of segregating sites in the alignment is low. Finally, HOME, which cannot consider multiple OTU strains per extant host, is not substantially affected by the sampling at random of a single OTU strain per host: its power is intermediate and its type-I error rate remains at 0% (Supplementary Fig. 14).

When simulating duplications and losses (or non-detection within hosts), we observed similar trends with an overall decrease in the power of all the approaches, an increase of the type-I error rate of ALE to 5%, and an increase of the type-I error rate of PACo under null model 1 (but not null model 2; Figure 2; Supplementary Figs. 15-17). Increasing the number of randomizations to 100,000 in our significance test does not fix this high type-I error, and more generally does not increase the performances of PACo (Supplementary Fig. 18).

### Empirical application

Rarefaction analyses on the gut microbiota of the 18 primate species revealed that the sequencing depth used in each sample was sufficient to saturate per-sample OTU richness, and that Shannon indices per species also reached a plateau when increasing the number of samples (Supplementary Fig. 19). We found a total of 149 95% OTUs, 86 97% OTUs, and 47 Swarm OTUs that are “core OTUs” present in more than 50% of the primate species. These core OTUs are on average detected in 12 host species, and represent a minor fraction of the total number of reads in the primate gut microbiota (28%, 14%, and 7%, respectively). The number of segregating sites and strains in their alignments is similar to those of the OTUs simulated using substitution rates ranging from μ=0.05 to μ=0.5 (Supplementary Figs. 1 & 20). The majority of these core OTUs present a significant cophylogenetic pattern when using global-fit approaches or ALE, corresponding to between 14% (based on 95% OTUs) and 2% (based on Swarm OTUs) of the total number of reads of the primate gut microbiota (Figure 3a). Conversely, HOME detected a significant cophylogenetic pattern in only 20% of the tested OTUs, corresponding to less than 7% of the total number of reads. The different approaches agreed on a small set of OTUs (10% of the core 95% OTUs) for which the cophylogenetic pattern is significant regardless of the approach used (Figure 3b). As expected, in OTUs without a cophylogenetic pattern, ALE inferred a lot of host-switches compared to codivergences (Supplementary Fig. 21), and HOME inferred high substitution rates and many host-switches (Supplementary Fig. 22). In OTUs with a significant cophylogenetic pattern, both methods inferred fewer, but a still significant number (∼5) of host-switches.

The much higher number of OTUs with a cophylogenetic pattern according to global-fit approaches and ALE compared to HOME could be linked to the higher type-I error rate of global-fit approaches and ALE, to the lower statistical power of HOME, or both. When we compared the number of segregating sites and hosts of the OTUs with or without a cophylogenetic pattern, we found that OTUs with a cophylogenetic pattern in global-fit approaches and ALE have less nucleotide variation and are present in a smaller number of hosts (Supplementary Fig. 19). The high type-I error rate of these approaches under these conditions (Supplementary Figs. 2-10) suggests that many of the OTUs for which they detected a cophylogenetic pattern are false positives. Conversely, OTUs with a cophylogenetic pattern according to HOME have more nucleotide variation and are present in a larger number of hosts. The gain of power of HOME with increased information (Supplementary Figs. 2-10) suggests that some vertically transmitted OTUs with little nucleotide variation or present in a few hosts were missed by the method. Hence, both the high type-I error rate of global-fit approaches and ALE and the low statistical power of HOME probably contribute to the contrasting results they provide.

When selecting several OTU strains per host species and applying global-fit approaches and ALE, more than 75% of the tested OTUs presented a significant cophylogenetic pattern (Supplementary Fig. 23). However, the type-I error rate of these approaches is higher when there are multiple strains per host species (especially PACo; Supplementary Figs. 12-17), suggesting that many of these OTUs are false positives.

Our test of geographically-driven cophylogenetic patterns between the Old World and the New World revealed that, at least for these data, HOME is more likely than the other approaches to identify OTUs that are truly vertically transmitted (Figure 4). Indeed, the majority of OTUs with a significant cophylogenetic pattern inferred with ParaFit, PACo, and ALE still had a significant cophylogenetic pattern when randomizing the associations between primate species and OTU strains within the Old and New Worlds. Conversely, the majority of OTUs with a significant cophylogenetic pattern inferred with HOME no longer had this cophylogenetic pattern when randomizing the associations between primate species and OTU strains based on geography, as expected if the pattern arises from vertical transmission rather than geographically structured pools of microbes. 24 out of the 149 ‘core’ 95% OTUs (*i*.*e*. 16% of them) presented a cophylogenetic pattern that did not arise from geographic structure according to HOME, corresponding to at most 5% of the total number of reads of the primate gut microbiota (Figure 4). We considered that these OTUs are likely to be vertically transmitted (but see Discussion). These vertically transmitted bacteria belonged mostly to the class Clostridia (phylum Firmicutes), especially the orders Lachnospirales and Oscillospirales, and to a lesser extent to the class Bacilli (phylum Firmicutes).

## Discussion

In this study, we used simulations to compare the statistical performances of different global-fit and event-based approaches to detect vertically transmitted OTUs in microbiota characterized by DNA metabarcoding. We found that the different approaches are rather complementary (Table 2). Their application to primate gut microbiota identifies vertically transmitted bacterial OTUs that represent a small fraction (∼5%) of the total number of reads.

### Pros and cons of different quantitative approaches to detect vertical transmission

The main advantage of global-fit methods is their computational efficiency. Depending on the size of the dataset in hand, there might be no other choice than to use these methods instead of event-based approaches. Global-fit methods generally have high statistical power. Also, they are robust to the presence of up to an intermediate number host-switches even though, unlike event-based approaches, they do not explicitly model these events. However, they also have an elevated type-I error rate when there are only a few segregating sites in the alignment. Although global-fit approaches typically use a randomization technique that independently randomizes which host species are associated with each OTU strain (“null model 1”), we found that shuffling the host species names instead (“null model 2”), as done in event-based approaches, reduces their type-I error. Null model 2 conserves the structure of the interactions, while null model 1 conserves only the number of host species associated with each OTU strain, which is less conservative. We therefore recommend using null model 2 in this context. Given that PACo tends to often have a higher type-I error than ParaFit and takes more time to run, we also recommend using ParaFit over PACo to detect vertically transmitted OTUs. Even when using ParaFit with null model 2, global-fit approaches have a higher type-I error rate than event-based ones. Although this would require further testing with simulations, our empirical analyses suggest that global-fit approaches have the highest difficulty to distinguish a geographic structure in the data from the signal of vertical transmission. These results suggest that event-based approaches should be preferred over global-fit ones when possible.

ALE outperforms all the other approaches on simulated data when OTUs have accumulated enough divergence, *i*.*e*. >10 segregating sites and/or >8 unique strains in the context of our simulations (Table 2). It not only has high power and a low type-I error rate, but it also accurately fits reconciliation events (host-switches, duplications, and losses) between the hosts and the trees of OTU strains. To evaluate the significance of the reconciliated scenarios, we recommend separately comparing the number of codivergences and host-switches against null expectations (as in Dorrell et al. 2021), rather than looking at the differences between the number of codivergences and host-switches (as in Groussin *et al*., 2017), as the latter strategy decreases the statistical power of the approach. ALE does not perform as well under situations with a low number of segregating sites. In this case, there is a lot of uncertainty in the reconstructed trees of OTU strains, and ALE is very slow to run. In addition, in this situation ALE has a higher type-I error rate when the OTU is not present in all host species (*i*.*e*. when there are losses), which is frequently the case in empirical data. We therefore do not recommend using ALE when the amount of variation in the OTU strains is too low. Also, while ALE has a much higher power than HOME and a low type-I error rate on simulated data with enough segregating sites, our results on empirical data suggest that ALE is in fact not conservative enough in its inference of vertical transmission. In particular, our empirical results suggest that ALE does not easily distinguish a geographic structure in the data from vertical transmission.

In contrast to the other approaches, HOME keeps a low type-I error rate, at least under all the situations we tested. The major drawback of HOME is that it has limited power. Another drawback is not handling multiple OTU strains per host species. In the presence of multiple strains, a user of HOME can randomly sample a single strain per host. However, as we showed in our simulations with within-host duplications, this contributes to further decreasing the statistical power of HOME. Therefore, HOME should be used only when within-host duplications are infrequent. On the positive side, in addition to the low type-I error rate, our empirical results suggest that HOME can often distinguish a geographic structure in the data from vertical transmission. This should ideally be tested further with simulations including heterogeneous pools of environmental microbes and phylogenetic signal in the host geographic distributions.

The distinct statistical performances of the different approaches can be explained by their constructions. Global-fit approaches do not model underlying processes, and therefore do not perform as well as event-based approaches. HOME seems to be less likely than global-fit approaches or ALE to infer vertical transmission for OTUs that present a strong geographical signal, which is likely due to the fact that HOME directly models DNA substitutions on the host phylogeny. For example, if the pools of OTU strains differ between the New and Old Worlds, they are unlikely to be particularly well modeled by a substitution process on the host tree, and the model therefore rejects the hypothesis of vertical transmission. Other processes than vertical transmission and geographic structure can generate a cophylogenetic pattern between host and trees of OTU strains (de Vienne et al., 2013), and statistical methods that are based on models that can represent these processes are more likely to perform better than those that do not.

We limited our analyses to the comparisons of four computational approaches that have previously been used for detecting vertical transmission in host-associated metabarcoding datasets. One of them (ALE) was originally developed for species-gene reconciliations purposes. Other approaches have been developed in this context, and could potentially also be useful for detecting vertical transmission (e.g. Bansal, Kellis, Kordi, & Kundu, 2018; Jacox, Chauve, Szöllosi, Ponty, & Scornavacca, 2016; Morel, Kozlov, Stamatakis, & Szöllősi, 2020), which could be explored in future simulation and empirical works. Meanwhile, we suggest simultaneously combining several approaches: ParaFit (and ALE when there are enough segregating sites) may be used to identify a larger set of potentially vertically transmitted OTUs, some of which might be false positives, while HOME may be used to identify a conservative set of vertically transmitted OTUs. The right number of vertically transmitted OTUs is likely included between both estimates.

In other non-bacterial systems, such as host-macroparasite systems, genetic data and species delineation for the parasites are generally of better quality than those obtained with metabarcoding data. We can use our comparison of statistical performances under simulations with high substitution to guide the choice of method to use in this case. For such systems, we recommend using ALE (with the Dorrell et al. (2021) criteria) when computationally feasible, and ParaFit (with null model 2) otherwise. When a cophylogenetic pattern is detected, we recommend carefully checking that this pattern is not linked to geographic structure in the data. Other event-based approaches that do not consider phylogenetic uncertainty and rely on maximum parsimony, *e*.*g*. eMPRess (Santichaivekin et al., 2021), are also likely to be valuable tools for detecting vertical transmission in such systems.

### Vertical transmission in the primate gut microbiota

We observed quantitative differences in the number of bacterial OTUs with a significant cophylogenetic pattern according to the different OTU clustering we performed. In particular, we detected >2 times fewer ‘core’ OTUs when using the Swarm clustering, resulting in >2 times fewer OTUs with a cophylogenetic pattern, maybe because this clustering method over-splits vertically transmitted bacteria that have accumulated too many divergences (Perez-Lamarque & Morlon, 2019). Using approaches that can handle multiple OTU strains per host species, we found many OTUs with a cophylogenetic pattern. These are likely false positives, given the high type-I error rate of these approaches in such conditions. These results suggest that when it is not clear whether multiple OTU strains correspond to real biological units and not PCR or sequencing error artifacts, it is preferable to simply pick the most abundant strain per host species and ignore duplication events.

Ideally, to assess whether cophylogenetic patterns were generated by vertical transmission, one also has to test whether the divergence times for the hosts match those of the OTU strains (de Vienne et al., 2013). We cannot robustly reconstruct the trees of OTU strains here, but we can examine the number of segregating sites, which range between 2 and 15 (within a single OTU) across bacterial OTUs from the primate gut. Given that the 16S rRNA gene diverges on average by 1% every 50 million years (Myr) (Ochman, Elwyn, & Moran, 1999), and that the primates are >65 Myr old, a metabarcoding marker with less than 10 segregating sites suggests divergence times for the OTU strains that match those of the hosts. Most of our alignments meet this criterium. When the number of segregating sites exceeds 10 (especially for 95% OTUs), alignments might either correspond to conglomerates of several vertically transmitted OTUs (Perez-Lamarque & Morlon, 2019) or to fast-evolving bacteria, like vertically transmitted bacteria with small population sizes (Moran, Munson, Baumann, & Ishikawa, 1993).

When removing OTUs whose cophylogenetic pattern arose from a phylogenetic signal in host geographic distribution, we estimated that less than 15% of the ‘core’ OTUs present in the gut microbiota of more than 50% of the primate species are vertically transmitted. These OTUs only represent a small fraction (∼5%) of the total number of bacterial reads. Accounting for the sometimes low statistical power of the approaches we used (<50% in some conditions), we may conclude that at most 30% of the ‘core’ OTUs in the bacterial gut microbiota of primates are vertically transmitted. Given that mammal gut microbiota can be composed of a large proportion of transient food-derived and/or environment-specific microbes that are unlikely to be faithfully vertically transmitted over more than 50 Myr (Amato et al., 2019; Nishida & Ochman, 2019), this estimate seems more realistic than larger ones. Among the bacteria inferred to be vertically transmitted, we found a large proportion in the order Clostridia (phylum Firmicutes), as found in previous analyses (Gaulke et al., 2018; Groussin et al., 2017; Perez-Lamarque & Morlon, 2019).

## Conclusion

Looking at vertically transmitted OTUs using metabarcoding datasets is challenging because of the low amount of information contained in metabarcoding marker genes. The different approaches that can be used for this purpose have complementary advantages and weaknesses. We recommend combining HOME, which has very infrequent false positives but limited power, with ALE (when there is enough variation in the alignments) or ParaFit, which have a higher power but many false positives. The ‘right’ number of vertically transmitted OTUs is likely between the estimates obtained with these approaches. We also recommend performing further checks, such as randomizing the host-bacteria associations within the main geographic areas of host distribution, in order to test whether the detected cophylogenetic patterns may have been generated by other processes than vertical transmissions. Applied to the gut microbiota of primates, we confirm that gut bacteria can be vertically transmitted, although most of the gut microbiota is not. Future work focusing on the specificities of these vertically transmitted bacteria would provide a better understanding of the mechanisms favoring vertical transmission for some particular bacterial lineages.

## Data Accessibility and Benefit-Sharing Section

### Data Accessibility Statement

ParaFit, PACo, and HOME are available as R functions. A tutorial on how to use HOME is available at https://github.com/BPerezLamarque/HOME/. Amended functions of ParaFit and PACo (from the R-packages ape (Paradis et al., 2004) and paco (Hutchinson et al., 2017)) are available at: https://github.com/BPerezLamarque/Scripts/tree/master/Comparing_methods_vertical_transmission/. ALE requires the installation of PhyloBayes and the software ALE (https://github.com/ssolo/ALE/) and is executable on a terminal; a tutorial is available at https://github.com/ssolo/ALE/#using-ale.

Both our scripts and simulations (DNA alignments of the OTUs) are publicly accessible through the Open Science Framework (osf) portal: osf.io/2rw36/. Raw data for the empirical analyses (from Amato el al. (2019)) are available at: https://www.ebi.ac.uk/ena/data/view/PRJEB22679.

### Benefits Generated

Benefits from this research accrue from the sharing of our data and results on public databases as described above.

## Author contributions

BPL and HM designed the study. BPL performed the analyses. BPL and HM wrote the manuscript.

## Acknowledgments

The authors acknowledge members of the BioDiv team at IBENS, V. Daubin, and B. Boussau for helpful discussions, as well as the Editor and three anonymous referees for their constructive comments. This work was supported by a doctoral fellowship from the École Normale Supérieure de Paris attributed to BPL. HM acknowledges support from the European Research Council (grant CoG-PANDA).

