## Supplementary for "Comparing different computational approaches for detecting long-term vertical transmission in host-associated microbiota"

### Supplementary Figures:

**Supplementary Figure 1: Large range of numbers of segregating sites and numbers of strains in the simulations:**
The numbers of segregating sites (a) and strains (*i.e.* haplotypes; b) are represented as a function of the simulated scenario: either strict vertical transmission (0 host-switch), vertical transmission with host-switches (5, 10, 15, or 20 switches), or independently evolving ("indep.").

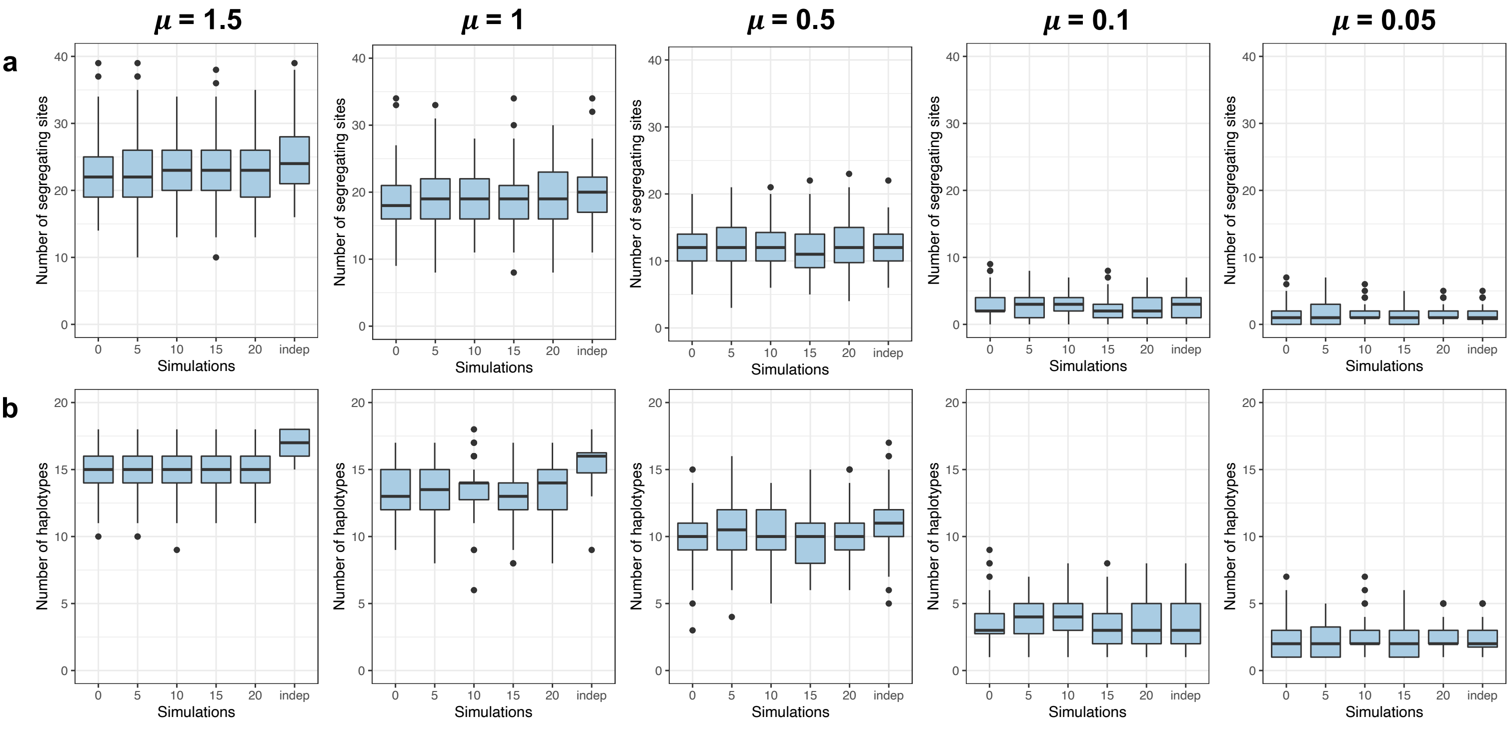

### Simulations without losses and duplications:

**Supplementary Figure 2: Computation time used by the different approaches as a function of the simulated substitution rates (μ).**

Boxplots indicate the computation time used by the different approaches for 10 OTU simulated with μ=1.5, μ=0.5, or μ=0.05. The simulations included different simulated scenarios: strict vertical transmission (0 host-switch) or independent evolution. Boxplots present the median surrounded by the first and third quartiles, and whiskers extend to the extreme values.

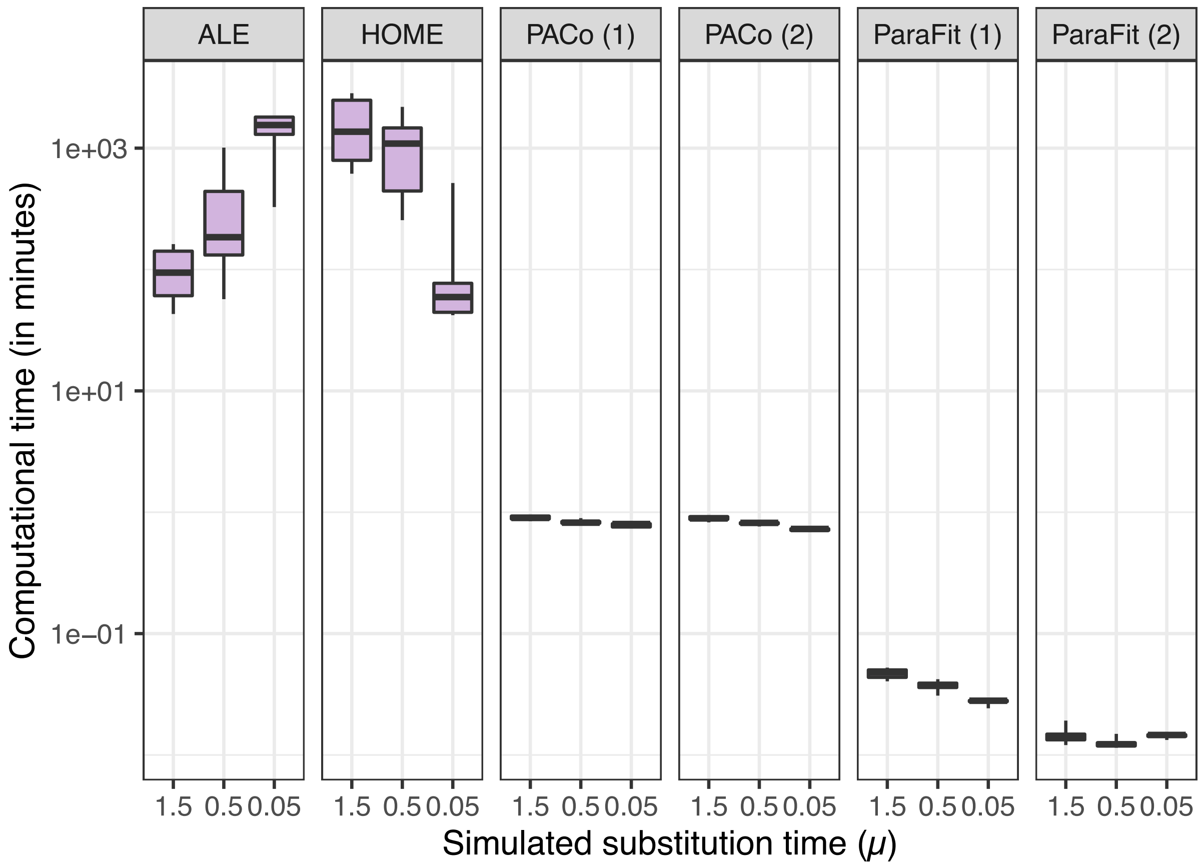

**Supplementary Figure 3: Statistical performances of the global-fit approach ParaFit evaluated based on the null model 1 (top panels) or the null model 2 (bottom panels) for different substitution rates (μ).**

Numbers of simulated OTUs rejecting the null hypothesis of independent evolution (rejected in red, not rejected in green, and not computed in grey) represented for different simulated scenarios: strict vertical transmission (0 host-switch), vertical transmission with host-switches (5, 10, 15, or 20 switches), and independent evolution ("indep."). Analyses measuring the statistical power and type-I error rate of the approach are plotted on a yellow and purple background, respectively. The average performance across scenarios is indicated as a percentage at the bottom of each panel.

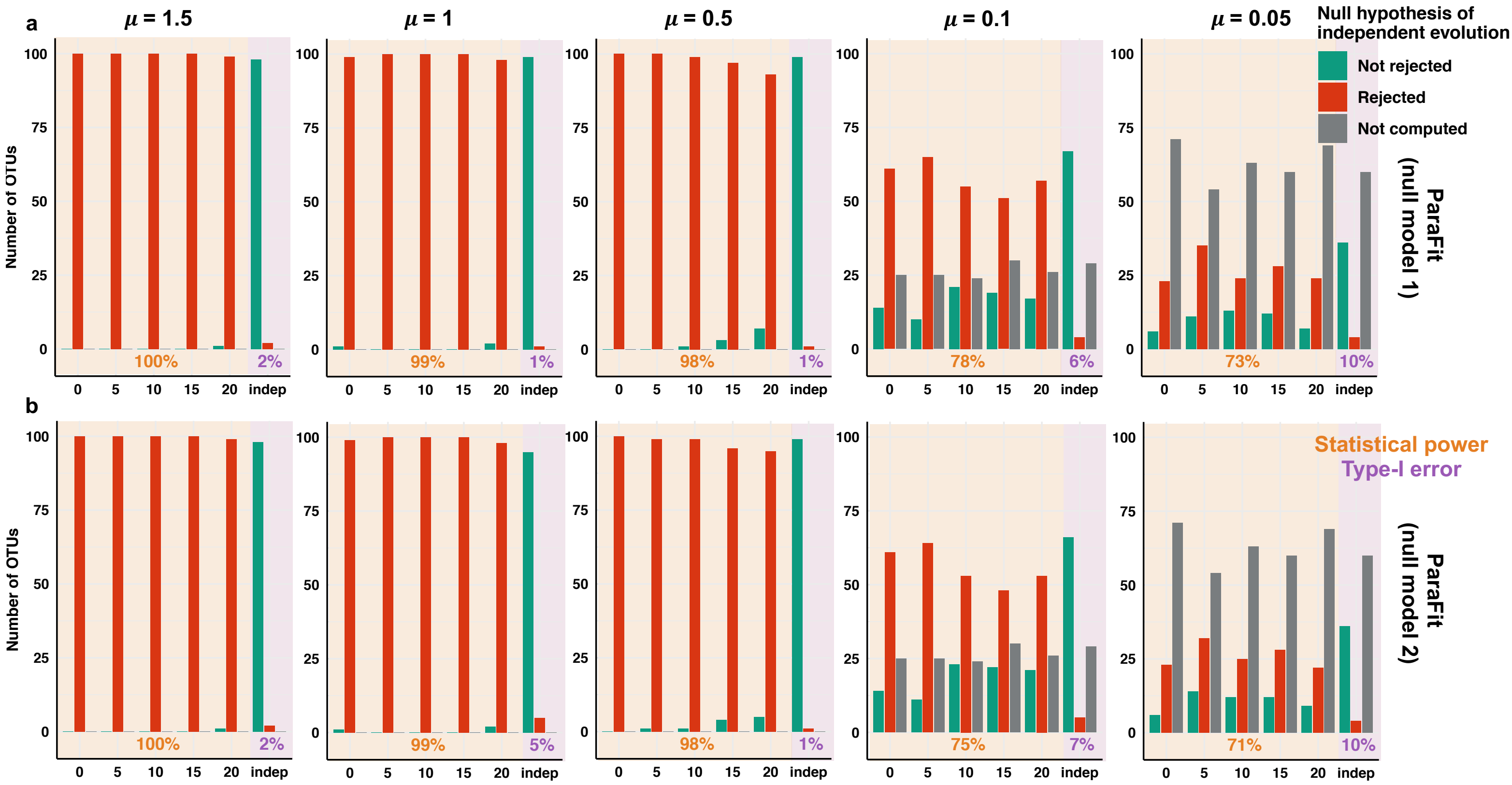

**Supplementary Figure 4: Statistical performances of the global-fit approach PACo evaluated based on the null model 1 (top panels) or the null model 2 (bottom panels) for different substitution rates (μ).**

Numbers of simulated OTUs rejecting the null hypothesis of independent evolution (rejected in red, not rejected in green, and not computed in grey) represented for different simulated scenarios: strict vertical transmission (0 host-switch), vertical transmission with host-switches (5, 10, 15, or 20 switches), and independent evolution ("indep."). Analyses measuring the statistical power and type-I error rate of the approach are plotted on a yellow and purple background, respectively. The average performance across scenarios is indicated as a percentage at the bottom of each panel.

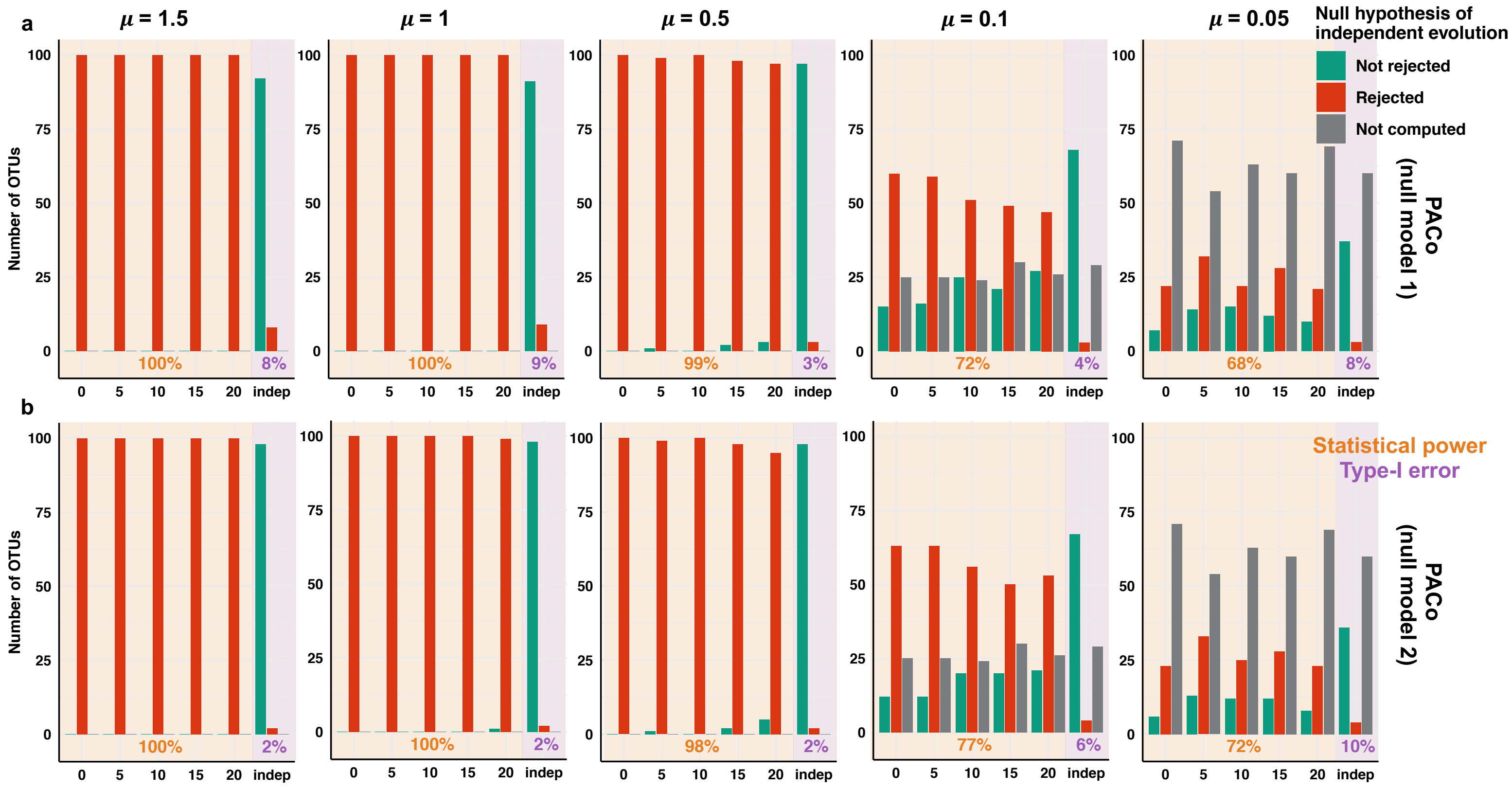

**Supplementary Figure 5: Statistical performances of the event-based approach ALE for different substitution rates (μ).**

(a) Numbers of simulated OTUs rejecting the null hypothesis of independent evolution (rejected in red, not rejected in green, and not computed in grey) represented for different simulated scenarios: strict vertical transmission (0 host-switch), vertical transmission with host-switches (5, 10, 15, or 20 switches), and independent evolution ("indep."). Rejections of the null hypothesis are performed using the criterium of Dorrell et al. (2021). Analyses measuring the statistical power and type-I error rate of the approach are plotted on a yellow and purple background, respectively. The average performance across scenarios is indicated as a percentage at the bottom of each panel. We did not run ALE for simulations with μ= 0.05, which were too long to compute.

(b) Estimated parameters (number of duplications, losses, codivergences, or host-switches) for μ=1.5 and μ=0.1 and different simulated scenarios: strict vertical transmission (0 host-switch), vertical transmission with 15 host-switches, and independent evolution ("indep."). Dark and light gray lines represent OTUs that are inferred to be transmitted or acquired from the environment, respectively.
Boxplots present the median surrounded by the first and third quartiles, and whiskers extend to the extreme values but no further than 1.5 of the inter-quartile range.

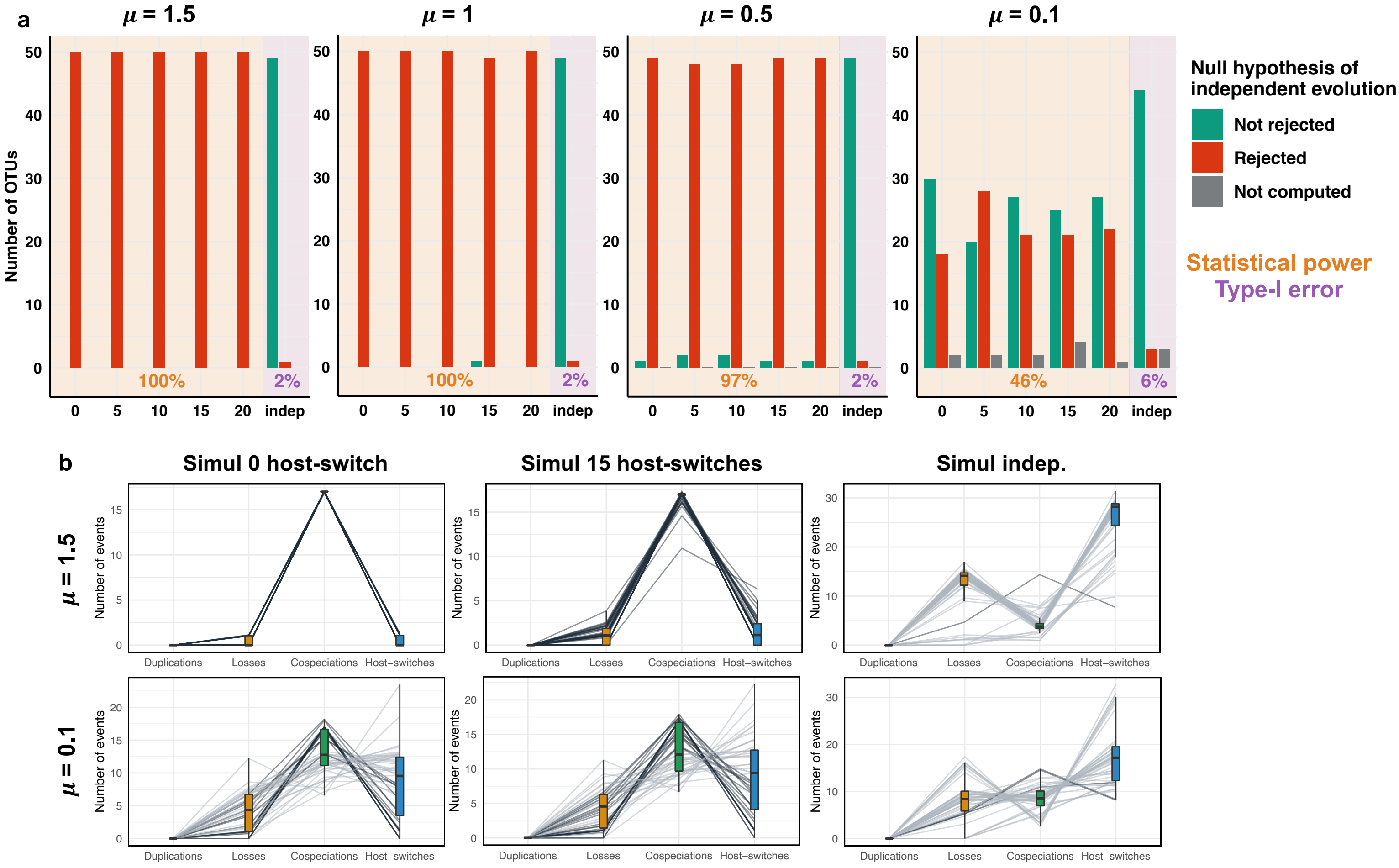

**Supplementary Figure 6: Statistical performances of the event-based approach ALE when using the criterium of Groussin et al. (2017) show low statistical power when substitution rates (μ) are low.**

Numbers of simulated OTUs rejecting the null hypothesis of independent evolution (rejected in red, not rejected in green, and not computed in grey) represented for different simulated scenarios: strict vertical transmission (0 host-switch), vertical transmission with host-switches (5, 10, 15, or 20 switches), and independent evolution ("indep."). Rejections of the null hypothesis are performed using the criterium of Groussin et al. (2017). Analyses measuring the statistical power and type-I error rate of the approach are plotted on a yellow and purple background, respectively. The average performance across scenarios is indicated as a percentage at the bottom of each panel. We did not run ALE for simulations with μ= 0.05, which were too long to compute.

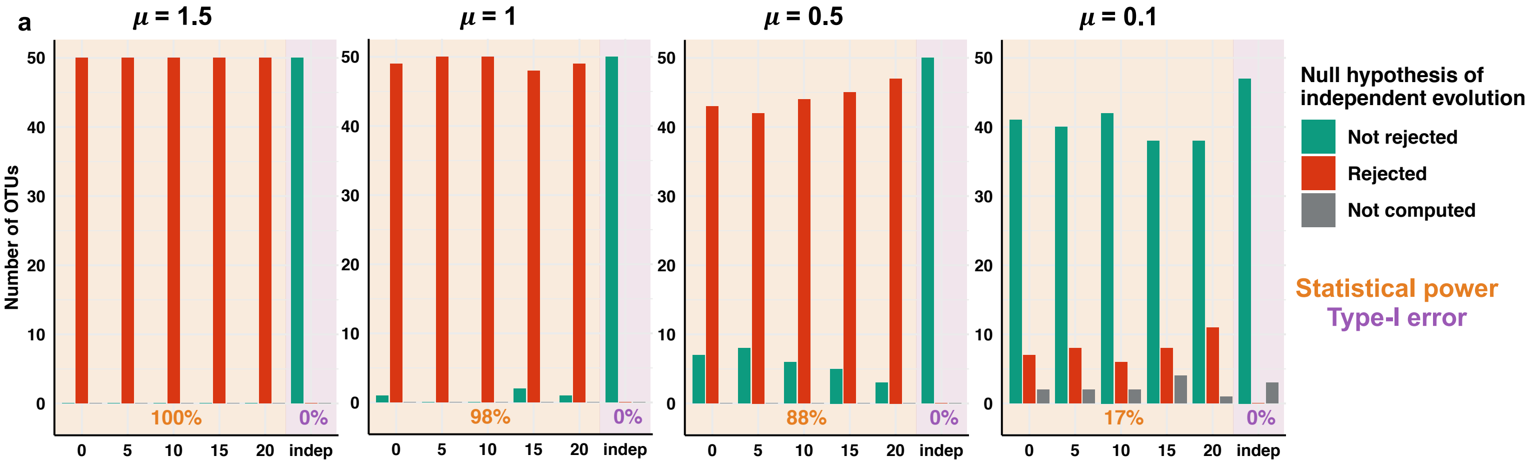

**Supplementary Figure 7: Statistical performances of the event-based approach HOME for different substitution rates (μ).**

(a) Number of simulated OTUs rejecting the null hypothesis of independent evolution (rejected in red, not rejected in green, and not computed in grey) represented for different simulated scenarios: either strict vertical transmission (0 host-switch), vertical transmission with host-switches (5, 10, 15, or 20 switches), or independent evolution ("indep."). Analyses measuring the statistical power and type-I error rate of the approach are plotted on a yellow and purple background, respectively. The average performance across scenarios is indicated as a percentage at the bottom of each panel.

(b) Estimated parameters (number of host-switches and substitution rates) for μ=1.5 and μ=0.1 and different simulated scenarios: strict vertical transmission (0 host-switch), vertical transmission with 15 host-switches, and independent evolution ("indep."). The "y" axes are square-root transformed. Dark and light gray lines represent OTUs that are inferred to be transmitted or acquired from the environment, respectively. Boxplots present the median surrounded by the first and third quartiles, and whiskers extend to the extreme values but no further than 1.5 of the inter-quartile range.

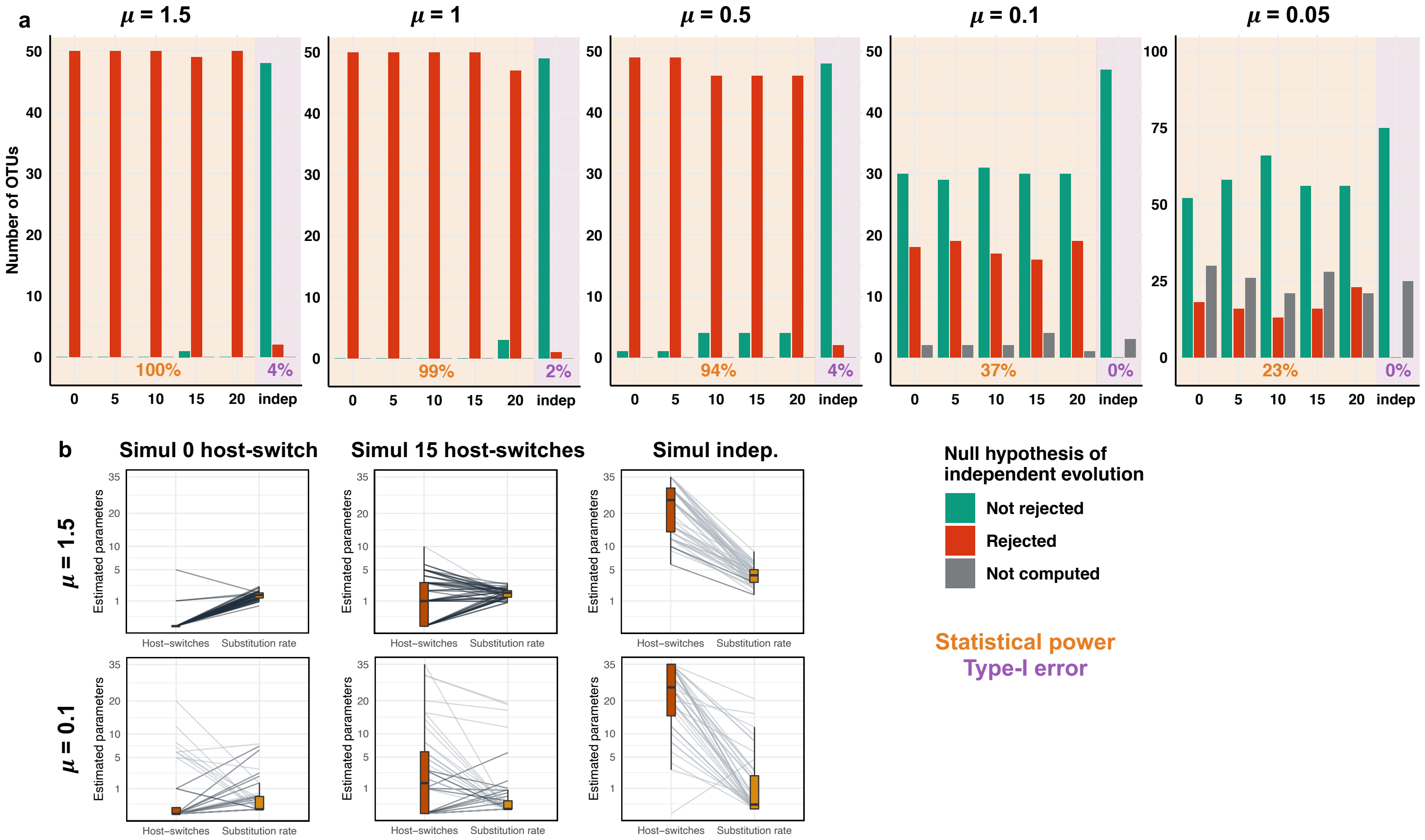

### Simulations with losses:

**Supplementary Figure 8: Statistical performances of the global-fit approaches, ParaFit (a) and PACo (b) when OTUs were simulated with losses (i.e. each OTU is only present in 10 host species).**

Numbers of simulated OTUs rejecting the null hypothesis of independent evolution (rejected in red, not rejected in green, and not computed in grey) represented as a function of the simulated scenario: either strict vertical transmission (0 host-switch), vertical transmission with host-switches (5, 10, 15, or 20 switches), or independently evolving ("indep.").

ParaFit and PACo were both evaluated using the null model 1 (top panels) and null model 2 (bottom panels).

Scenarios showing the statistical power of the approach are highlighted in yellow, whereas the ones indicating the type-I error rate are in purple: these performances are indicated as percentages at the bottom of the panels.

Each panel corresponds to the different simulated substitution rates (μ).

Although global-fit approaches appeared to have an important statistical power when μ=0.05, note that they were not able to run for ~75% of the simulated OTUs (because they had less than 3 strains).

(a) ParaFit

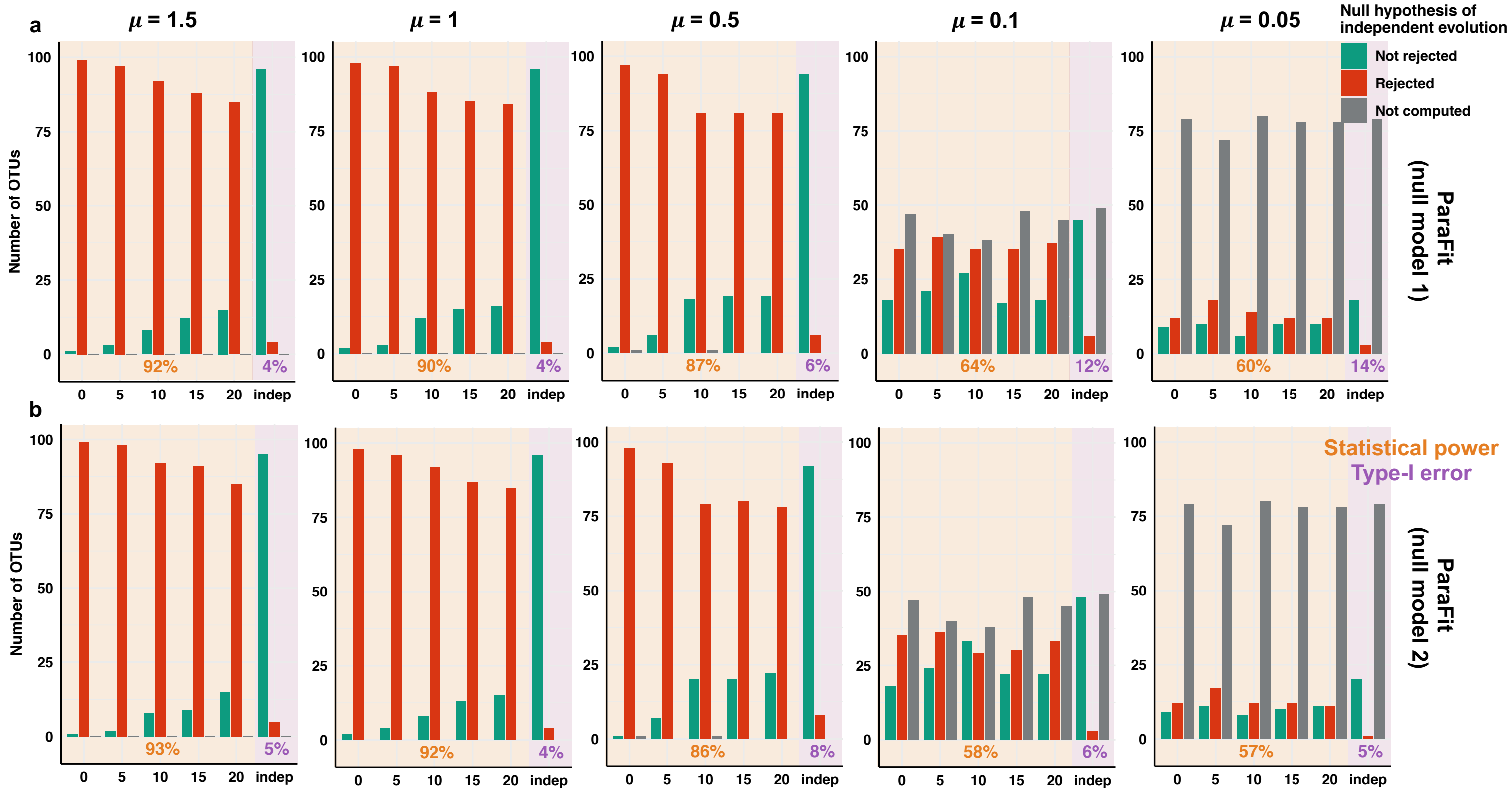

(b) PACo

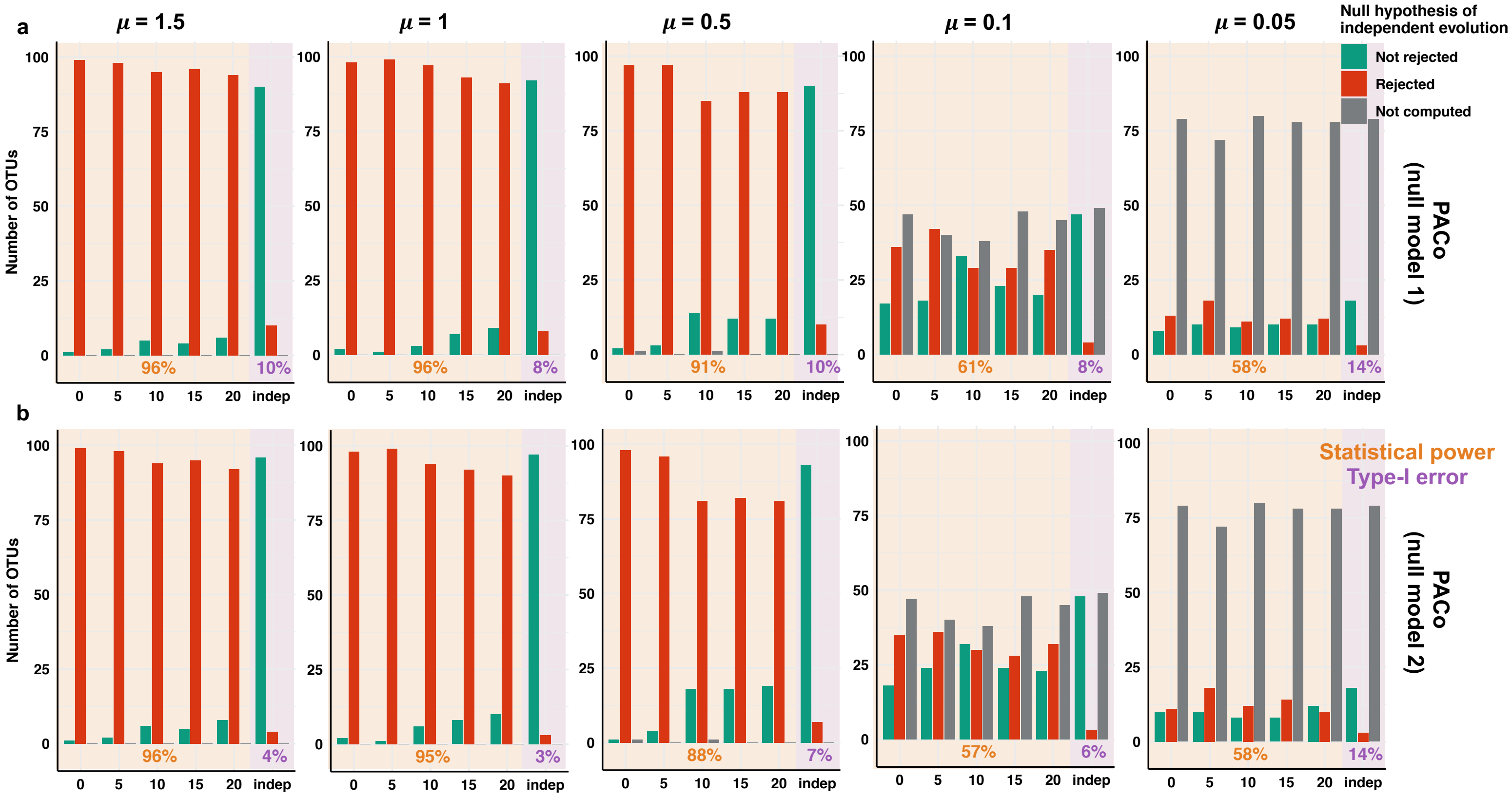

a
μ = 1.5
μ = 1
μ = 0.5
μ = 0.1
μ = 0.05
Null hypothesis of independent evolution
Not rejected
Rejected
Not computed
Number of OTUs
96%
10%
96%
8%
91%
10%
61%
8%
58%
14%
0 5 10 15 20 indep
PACo (null model 1)
b
Statistical power
Type-I error
96%
4%
95%
3%
88%
7%
57%
6%
58%
14%
PACo (null model 2)

**Supplementary Figure 9: Statistical performances of ALE when OTUs were simulated with losses (i.e. each OTU is only present in 10 host species).**

Numbers of simulated OTUs rejecting the null hypothesis of independent evolution (rejected in red, not rejected in green, and not computed in grey) represented as a function of the simulated scenario: either strict vertical transmission (0 host-switch), vertical transmission with host-switches (5, 10, 15, or 20 switches), or independently evolving (“indep.”).

Scenarios showing the statistical power of the approach are highlighted in yellow, whereas the ones indicating the type-I error rate are in purple: these performances are indicated as percentages at the bottom of the panels.

Each panel corresponds to the different simulated substitution rates (μ), except μ= 0.05, which was too long to be computed.

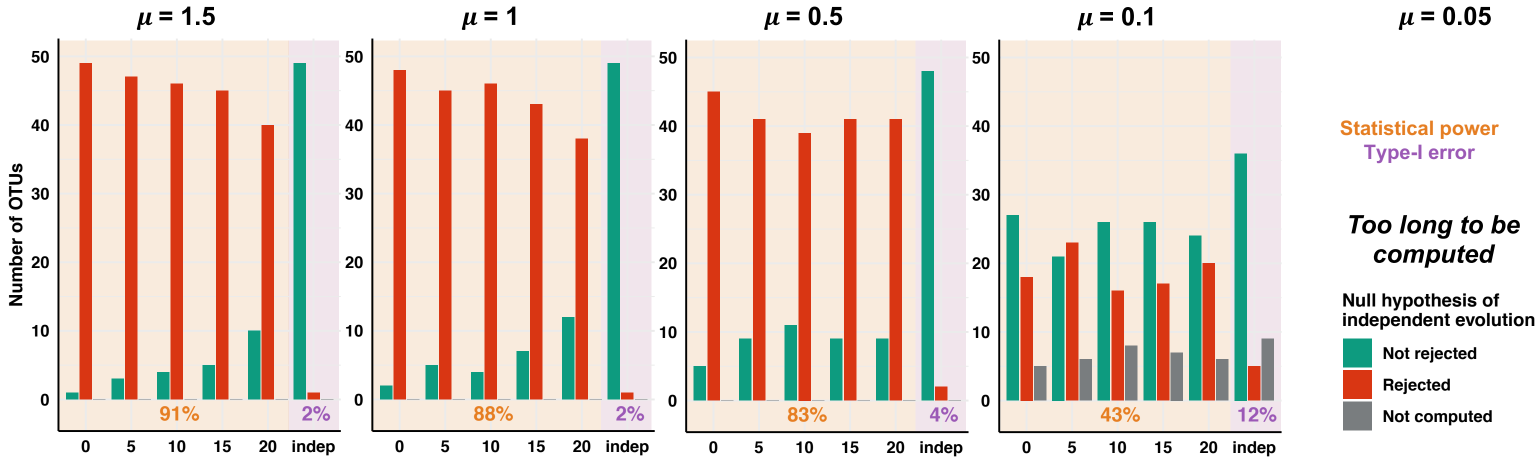

**Supplementary Figure 10: Statistical performances of HOME when OTUs were simulated with losses (i.e. each OTU is only present in 10 host species).**

Numbers of simulated OTUs rejecting the null hypothesis of independent evolution (rejected in red, not rejected in green, and not computed in grey) represented as a function of the simulated scenario: either strict vertical transmission (0 host-switch), vertical transmission with host-switches (5, 10, 15, or 20 switches), or independently evolving ("indep.").

Scenarios showing the statistical power of the approach are highlighted in yellow, whereas the ones indicating the type-I error rate are in purple: these performances are indicated as percentages at the bottom of the panels.

Each panel corresponds to the different simulated substitution rates (μ).

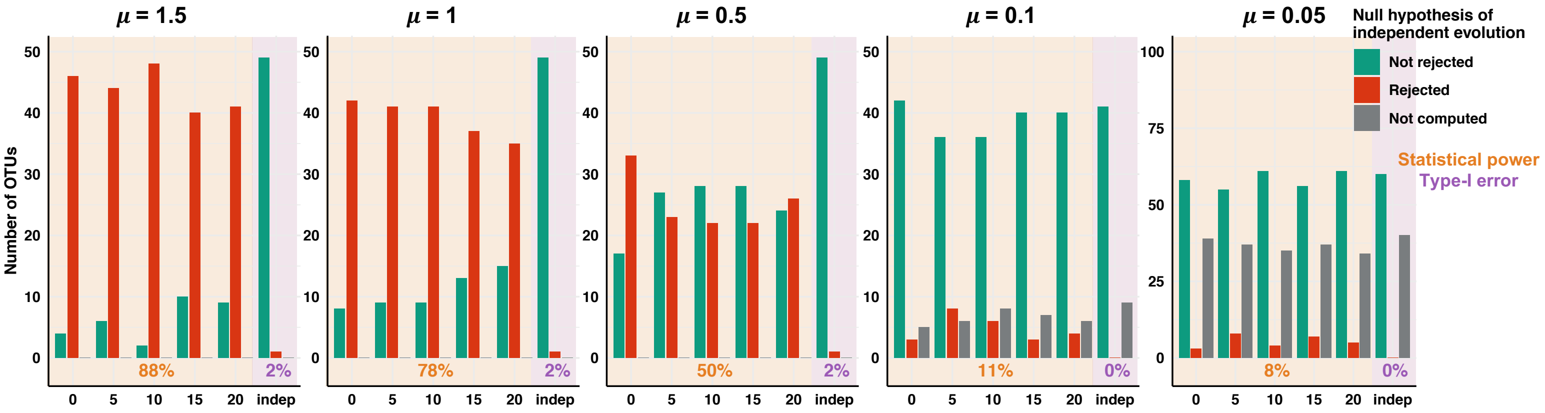

### Simulations with duplications:

**Supplementary Figure 11: Large range of numbers of segregating sites and numbers of strains in the simulations with duplications:**

The numbers of segregating sites (a) and strains (i.e. haplotypes; b) are represented as a function of the simulated scenario: either strict vertical transmission (0 host-switch), vertical transmission with host-switches (5, 10, 15, or 20 switches), or independently evolving (“indep.”).

Boxplots present the median surrounded by the first and third quartiles, and whiskers extend to the extreme values but no further than 1.5 of the inter-quartile range.

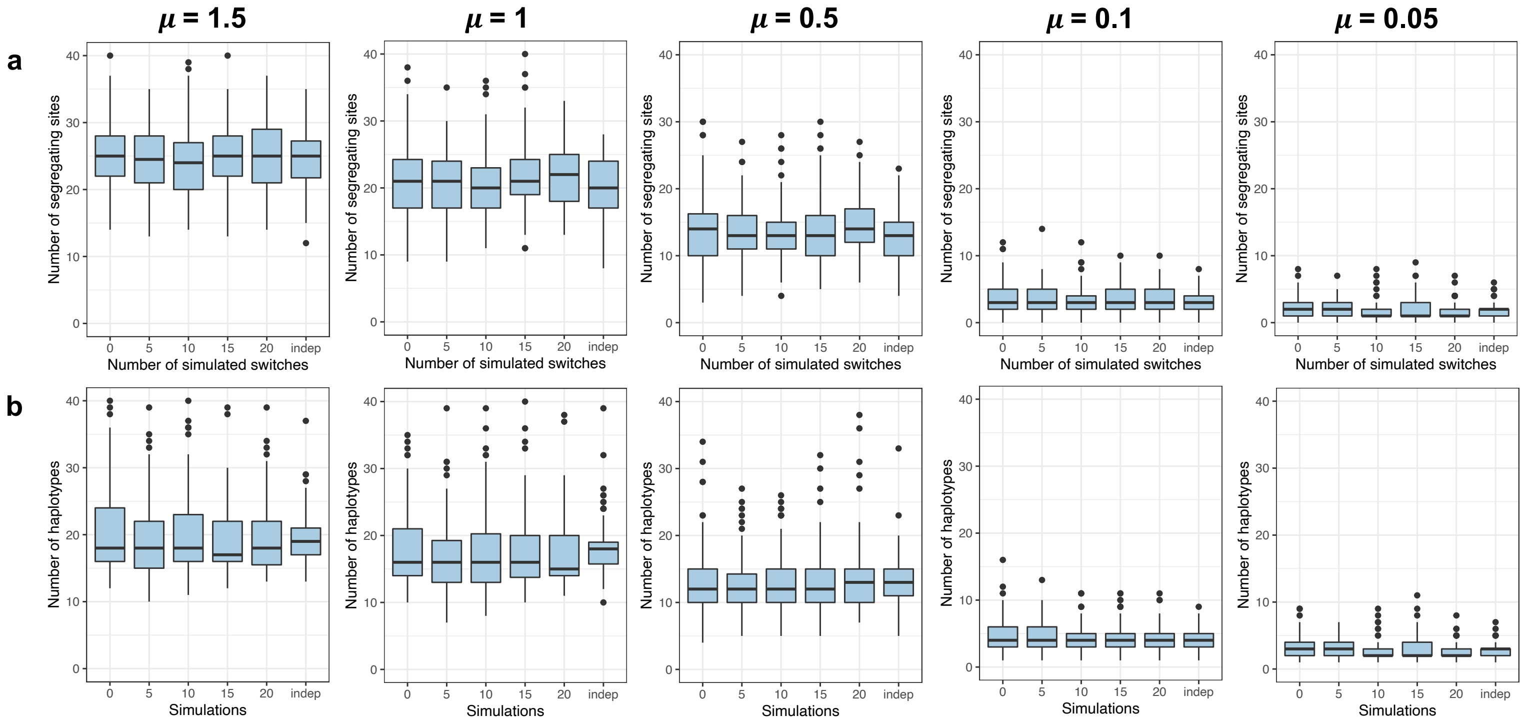

**Supplementary Figure 12: Statistical performances of the global-fit approaches, ParaFit (a) and PACo (b) when simulating duplications.**

Numbers of simulated OTUs rejecting the null hypothesis of independent evolution (rejected in red, not rejected in green, and not computed in grey) represented as a function of the simulated scenario: either strict vertical transmission (0 host-switch), vertical transmission with host-switches (5, 10, 15, or 20 switches), or independently evolving ("indep.").

ParaFit and PACo were both evaluated using the null model 1 (top panels) and null model 2 (bottom panels).

Scenarios showing the statistical power of the approach are highlighted in yellow, whereas the ones indicating the type-I error rate are in purple: these performances are indicated as percentages at the bottom of the panels.

Each panel corresponds to the different simulated substitution rates (μ).

(a) ParaFit

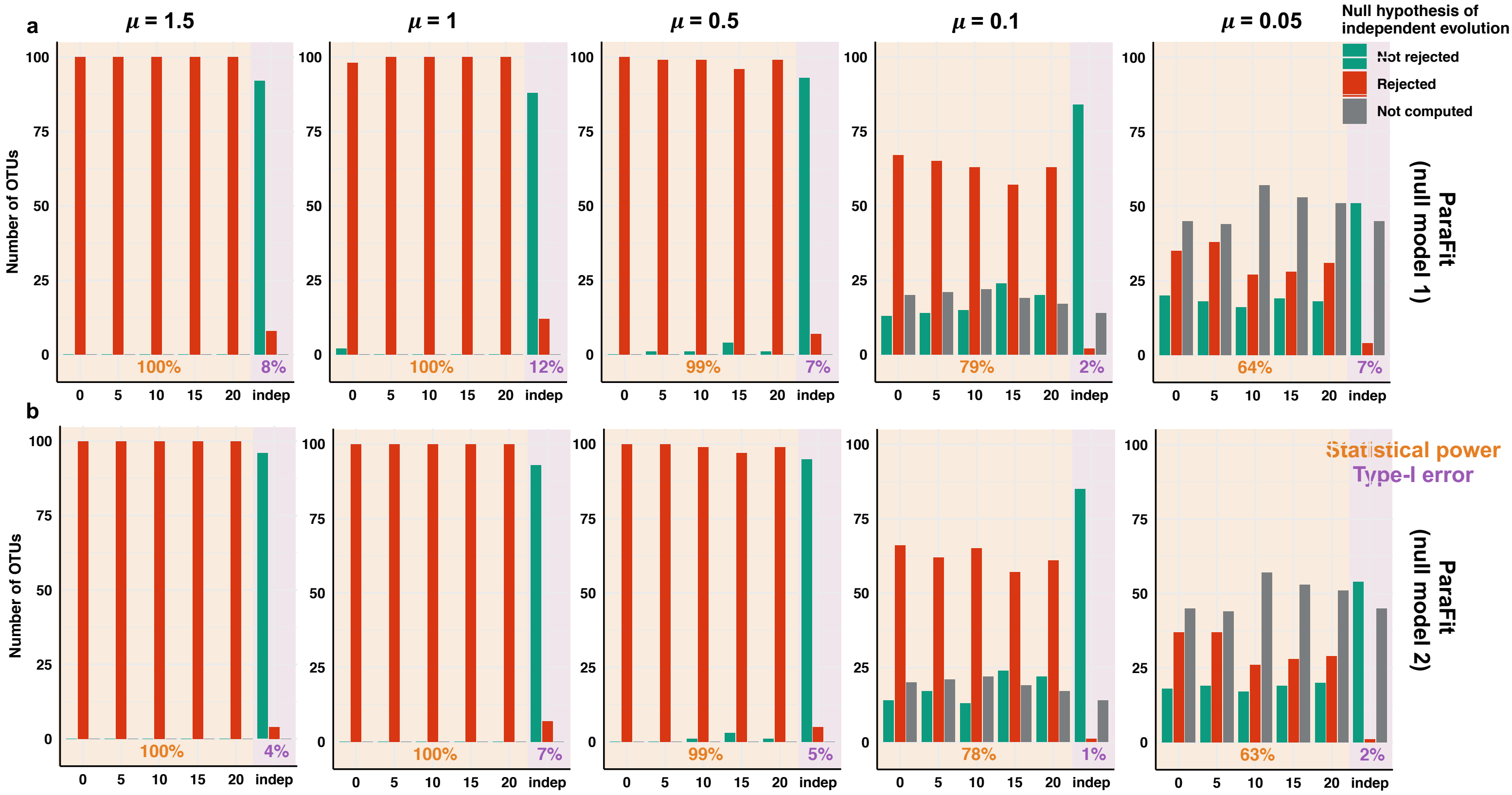

(b) PACo

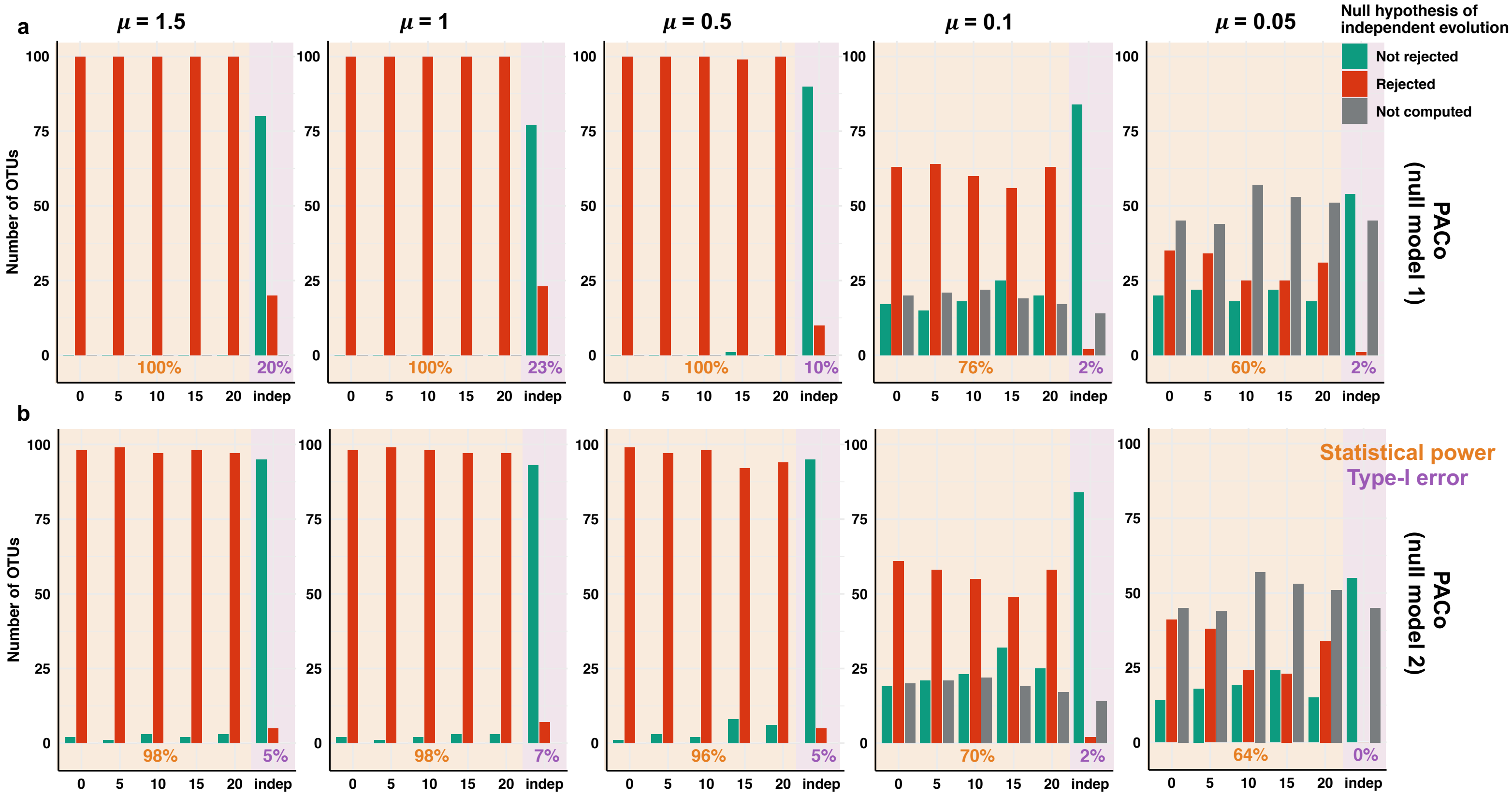
a
b
μ = 1.5
μ = 1
μ = 0.5
μ = 0.1
μ = 0.05
Number of OTUs
Null hypothesis of independent evolution
Not rejected
Rejected
Not computed
PACo (null model 1)
PACo (null model 2)
Statistical power
Type-I error
100%
20%
100%
23%
100%
10%
76%
2%
60%
2%
98%
5%
98%
7%
96%
5%
70%
2%
64%
0%
0
5
10
15
20
indep

**Supplementary Figure 13: Statistical performances of ALE when simulating duplications.**

Numbers of simulated OTUs rejecting the null hypothesis of independent evolution (rejected in red, not rejected in green, and not computed in grey) represented as a function of the simulated scenario: either strict vertical transmission (0 host-switch), vertical transmission with host-switches (5, 10, 15, or 20 switches), or independently evolving (“indep.”).

Scenarios showing the statistical power of the approach are highlighted in yellow, whereas the ones indicating the type-I error rate are in purple: these performances are indicated as percentages at the bottom of the panels.

Each panel corresponds to the different simulated substitution rates (μ), except μ= 0.05, which was too long to be computed.

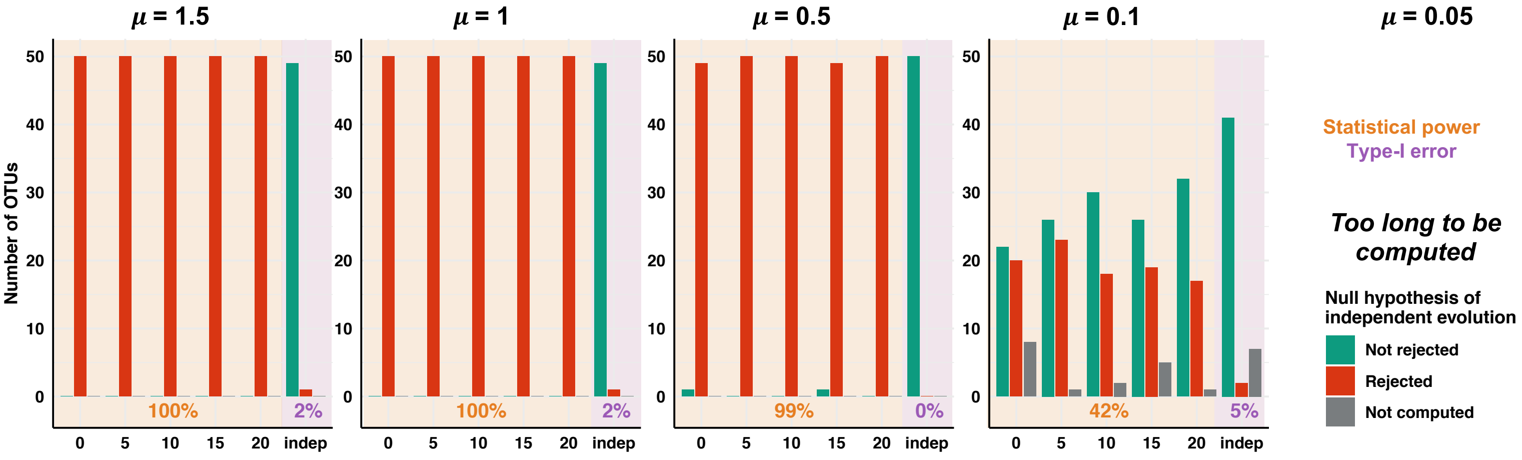

**Supplementary Figure 14: Statistical performances of HOME when simulating duplications.**

Numbers of simulated OTUs rejecting the null hypothesis of independent evolution (rejected in red, not rejected in green, and not computed in grey) represented as a function of the simulated scenario: either strict vertical transmission (0 host-switch), vertical transmission with host-switches (5, 10, 15, or 20 switches), or independently evolving (“indep.”).

Scenarios showing the statistical power of the approach are highlighted in yellow, whereas the ones indicating the type-I error rate are in purple: these performances are indicated as percentages at the bottom of the panels: these performances are indicated as percentages at the bottom of the panels.

Each panel corresponds to the different simulated substitution rates (μ), except μ=1.5 or μ=1, which were too long to be computed.

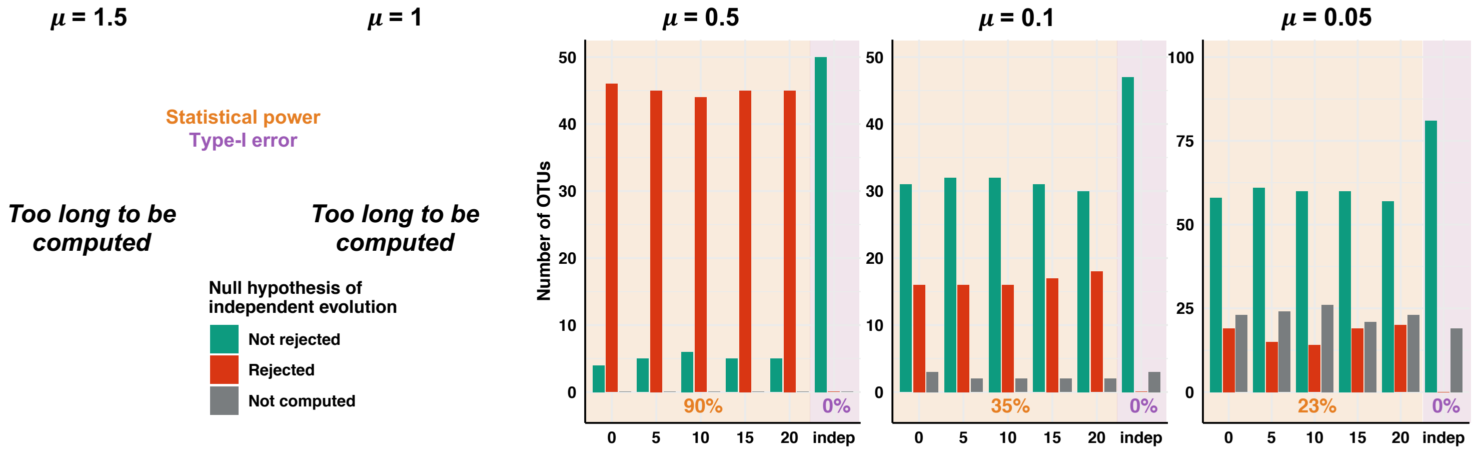

### Simulations with duplications and losses:

**Supplementary Figure 15: Statistical performances of the global-fit approaches, ParaFit (a) and PACo (b) when OTUs were simulated with duplications and losses (i.e. each OTU is only present in 10 host species).**
Numbers of simulated OTUs rejecting the null hypothesis of independent evolution (rejected in red, not rejected in green, and not computed in grey) represented as a function of the simulated scenario: either strict vertical transmission (0 host-switch), vertical transmission with host-switches (5, 10, 15, or 20 switches), or independently evolving ("indep.").
ParaFit and PACo were both evaluated using the null model 1 (top panels) and null model 2 (bottom panels).
Scenarios showing the statistical power of the approach are highlighted in yellow, whereas the ones indicating the type-I error rate are in purple: these performances are indicated as percentages at the bottom of the panels.
Each panel corresponds to the different simulated substitution rates (μ).

(a) ParaFit

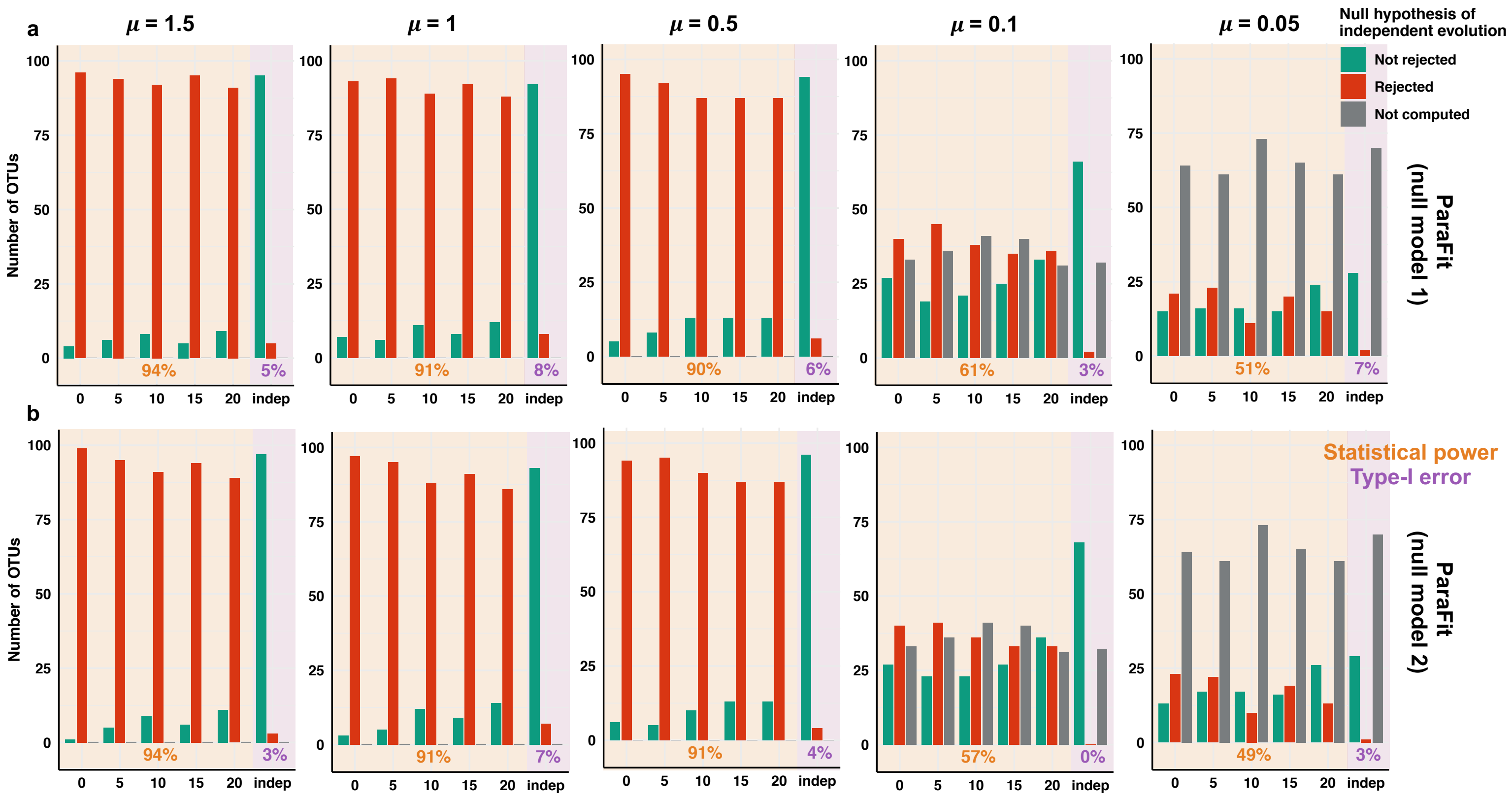

(b) PACo

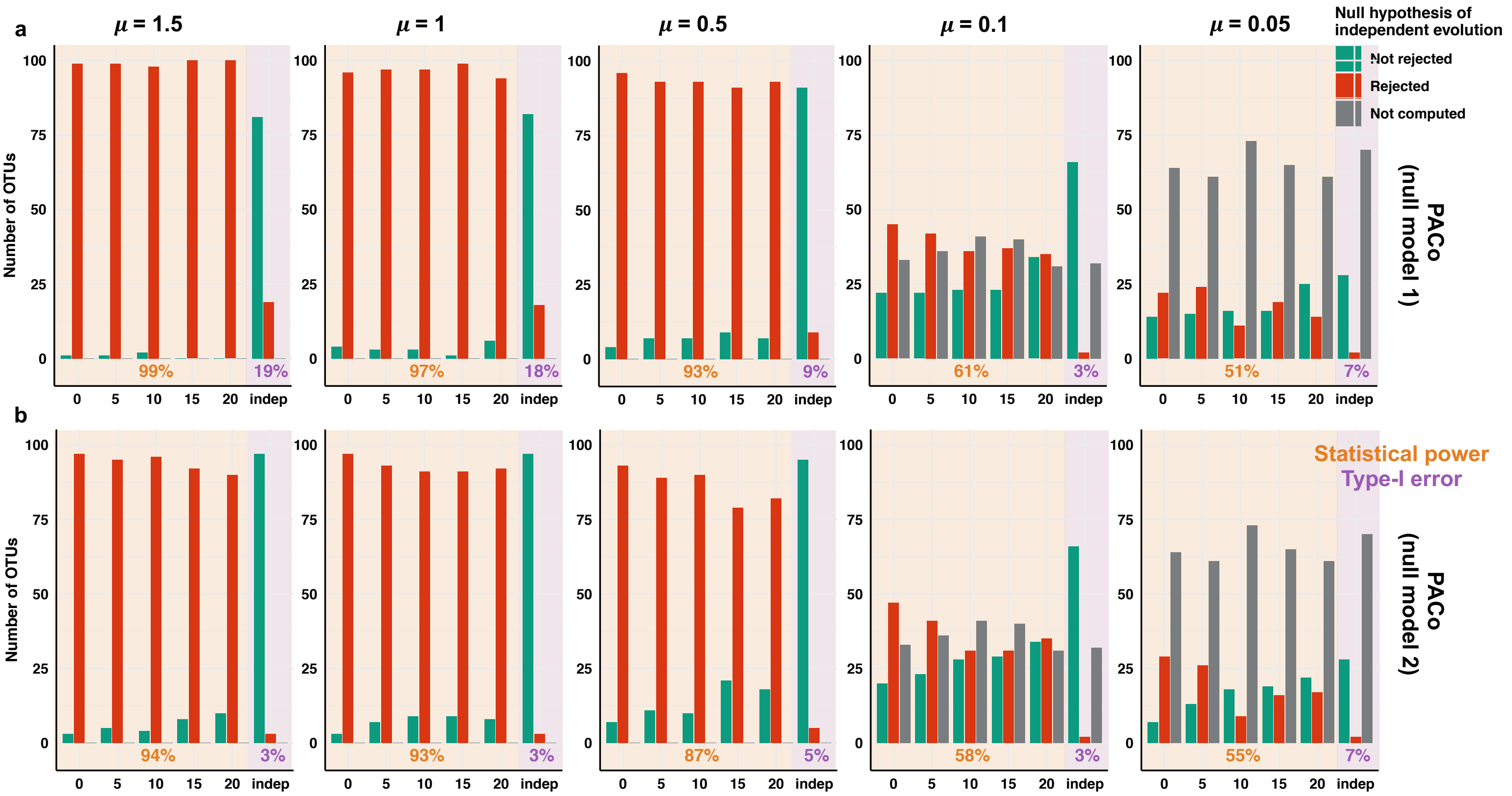

a
μ = 1.5
μ = 1
μ = 0.5
μ = 0.1
μ = 0.05
Null hypothesis of independent evolution
Not rejected
Rejected
Not computed
Number of OTUs
PACo (null model 1)
99%
19%
97%
18%
93%
9%
61%
3%
51%
7%
b
Statistical power
Type-I error
PACo (null model 2)
94%
3%
93%
3%
87%
5%
58%
3%
55%
7%
0 5 10 15 20 indep

**Supplementary Figure 16: Statistical performances of ALE when OTUs were simulated with duplications and losses (i.e. each OTU is only present in 10 host species).**

Numbers of simulated OTUs rejecting the null hypothesis of independent evolution (rejected in red, not rejected in green, and not computed in grey) represented as a function of the simulated scenario: either strict vertical transmission (0 host-switch), vertical transmission with host-switches (5, 10, 15, or 20 switches), or independently evolving ("indep.").

Scenarios showing the statistical power of the approach are highlighted in yellow, whereas the ones indicating the type-I error rate are in purple: these performances are indicated as percentages at the bottom of the panels.

Each panel corresponds to the different simulated substitution rates (μ), except μ= 0.05, which was too long to be computed.

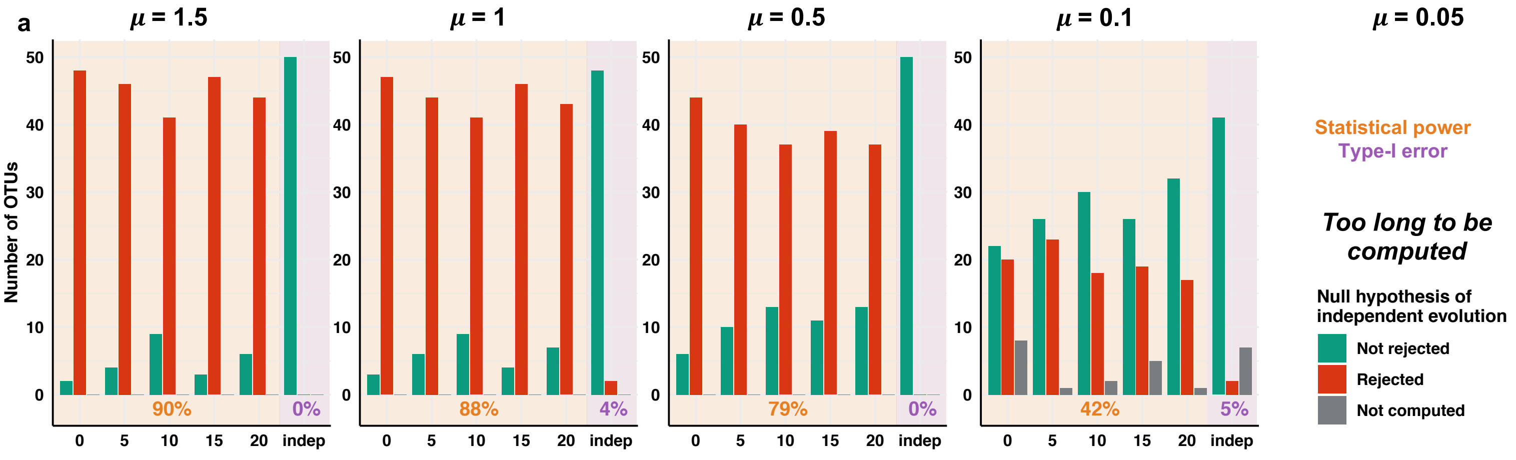

**Supplementary Figure 17: Statistical performances of HOME when OTUs were simulated with duplications and losses (i.e. each OTU is only present in 10 host species).**

Numbers of simulated OTUs rejecting the null hypothesis of independent evolution (rejected in red, not rejected in green, and not computed in grey) represented as a function of the simulated scenario: either strict vertical transmission (0 host-switch), vertical transmission with host-switches (5, 10, 15, or 20 switches), or independently evolving ("indep.").

Scenarios showing the statistical power of the approach are highlighted in yellow, whereas the ones indicating the type-I error rate are in purple: these performances are indicated as percentages at the bottom of the panels.

Each panel corresponds to the different simulated substitution rates (μ), except μ=1.5 or μ=1, which were too long to be computed.

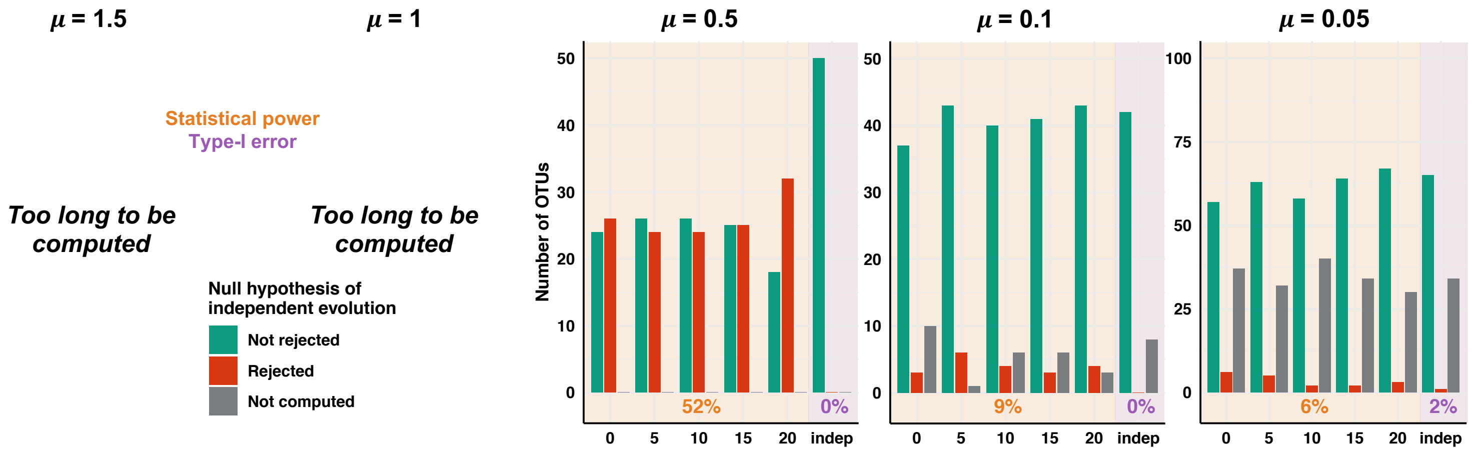

**Supplementary Figure 18: Using 100,000 permutations instead of 10,000 did not significantly affect the statistical performances of PACo.**

Numbers of simulated OTUs (simulated with duplications and losses) rejecting the null hypothesis of independent evolution (rejected in red, not rejected in green, and not computed in grey) represented for different simulated scenarios: strict vertical transmission (0 host-switch), vertical transmission with host-switches (5, 10, 15, or 20 switches), and independent evolution ("indep."). Analyses measuring the statistical power and type-I error rate of the approach are plotted on a yellow and purple background, respectively. The average performance across scenarios is indicated as a percentage at the bottom of each panel. We report here results obtained with null model 1 (null model 2 gave very similar results).

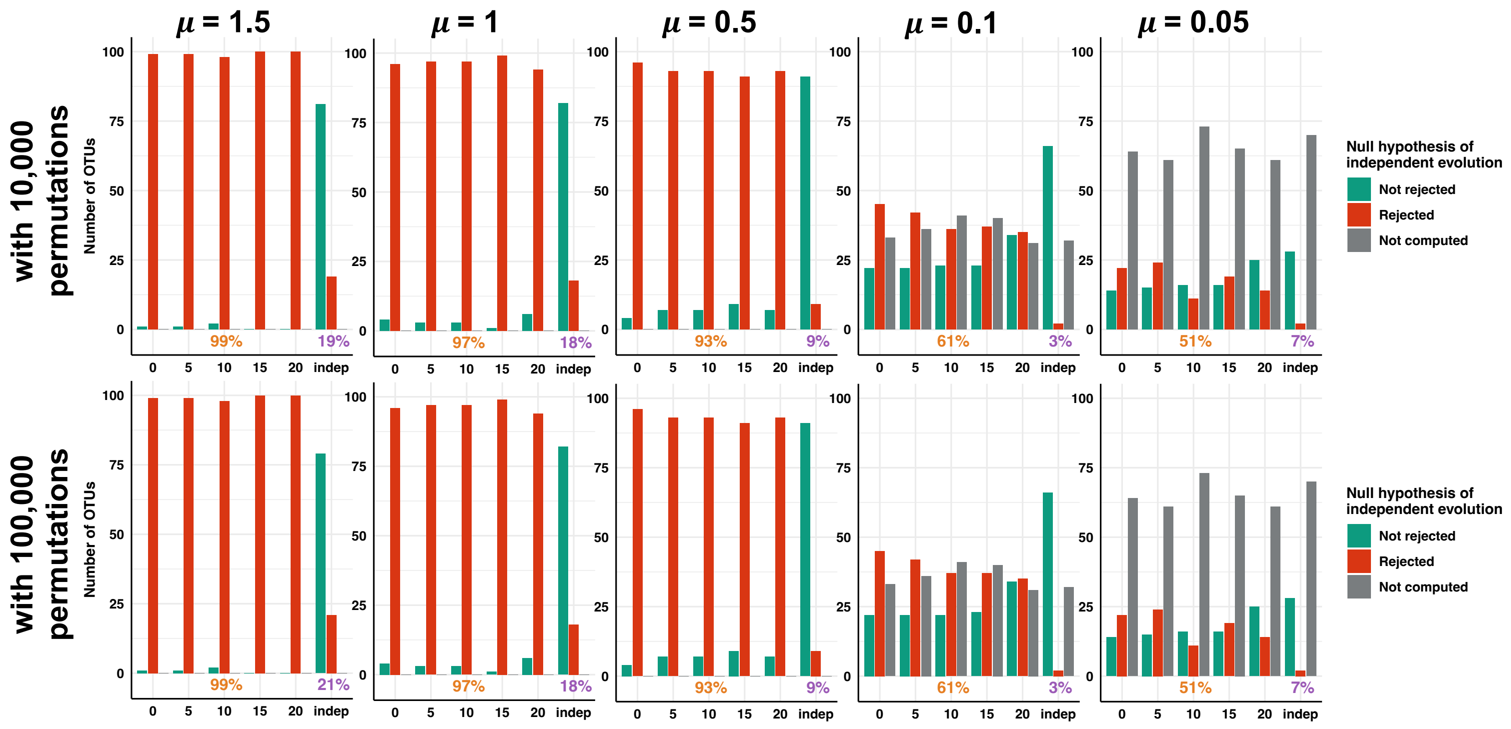

### Empirical application:

**Supplementary Figure 19: Rarefactions analyses indicated that most of the bacterial diversity is documented in each sample (a) and in each species (b).**

**(a)** Rarefaction curves: Number of OTU (OTU richness) as a function of the number of reads per sample. Rarefaction were repeated 10 times per number of subsampled reads.

**(b)** Shannon diversity as a function of the number of gut microbiota samples per primate species. For each species, we draw rarefaction curves by subsampling a given number of samples corresponding to this species and computed the associated Shannon diversity. For each number of species, we replicated 100 times the subsampling.

OTUs were either clustered as 95%, 97%, or as Swarm OTUs.

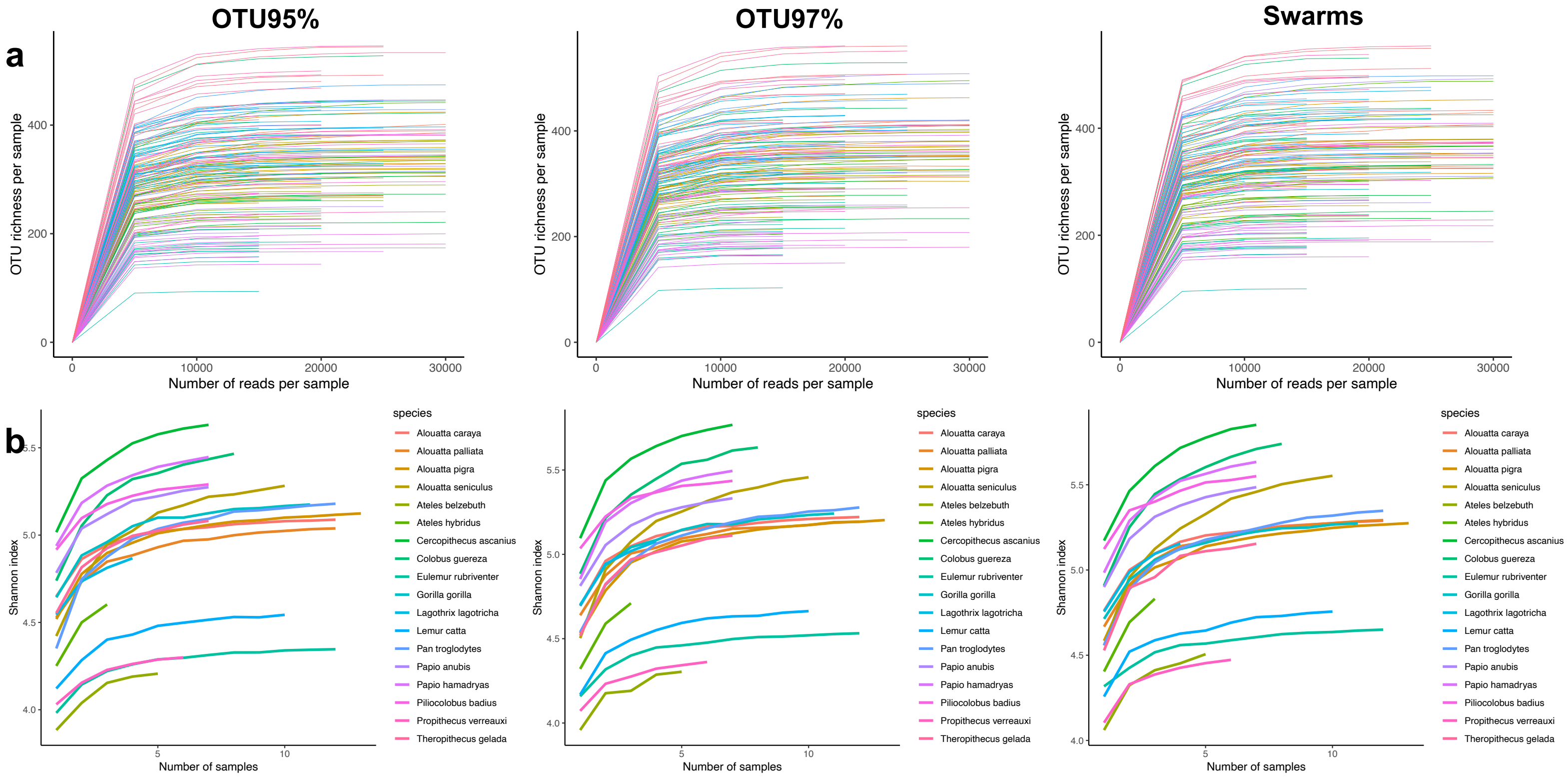

**Supplementary Figure 20: Large range of numbers of segregating sites and numbers of strains in the bacterial OTUs of the primate gut microbiota**

For each OTU, the numbers of segregating sites (a), the numbers of host species where the OTU was found (b), and the numbers of strains (i.e. haplotypes; c) are represented as a function of the simulated scenario: either strict vertical transmission (0 host-switch), vertical transmission with host-switches (5, 10, 15, or 20 switches), or independently evolving ("indep."). OTUs were either clustered as 95%, 97%, or as Swarm OTUs.

All the OTUs are represented on the left panels, and then on the four right panels, we separated the OTUs according to their rejection or not of the null hypothesis of independent evolutions based on ALE, HOME, ParaFit, or ALE. ParaFit and PACo were both evaluated using the null model 1 (top panels) and null model 2 (bottom panels).

Boxplots present the median surrounded by the first and third quartiles, and whiskers extend to the extreme values but no further than 1.5 of the inter-quartile range.

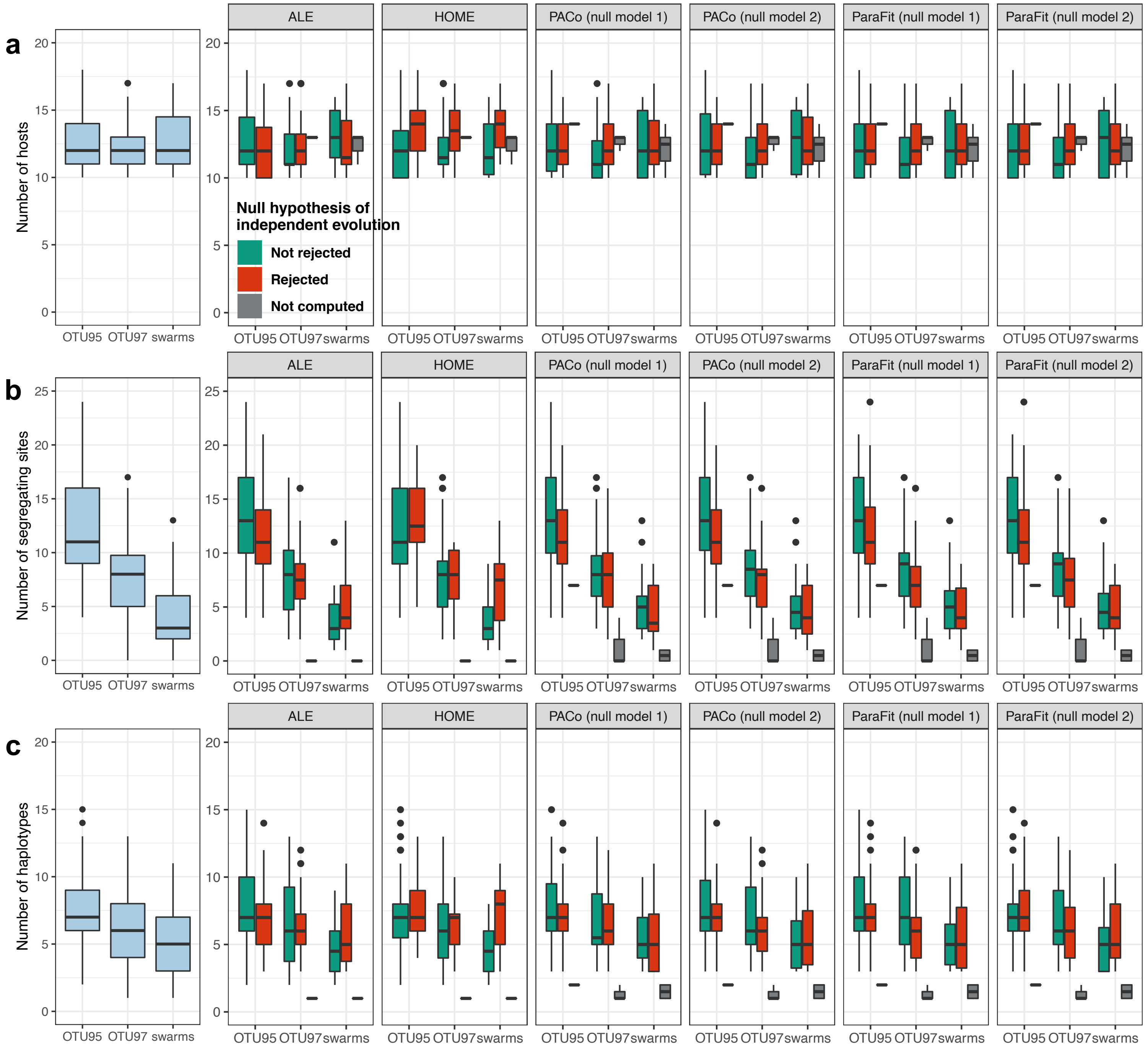
a
b
c
Number of hosts
Number of segregating sites
Number of haplotypes
ALE
HOME
PACo (null model 1)
PACo (null model 2)
ParaFit (null model 1)
ParaFit (null model 2)
Null hypothesis of independent evolution
Not rejected
Rejected
Not computed
OTU95
OTU97
swarms

**Supplementary Figure 21: Number of reconciliation events estimated using ALE for the primate gut microbiota bacterial OTUs**

Estimated number of duplications, losses, codivergences, or host-switches for various OTU clustering criteria (either 95%, 97% sequence similarity, or Swarm OTUs), for the OTUs rejecting the null hypothesis of independent evolution (left) or not rejecting it (right).

Boxplots present the median surrounded by the first and third quartiles, and whiskers extend to the extreme values but no further than 1.5 of the inter-quartile range.

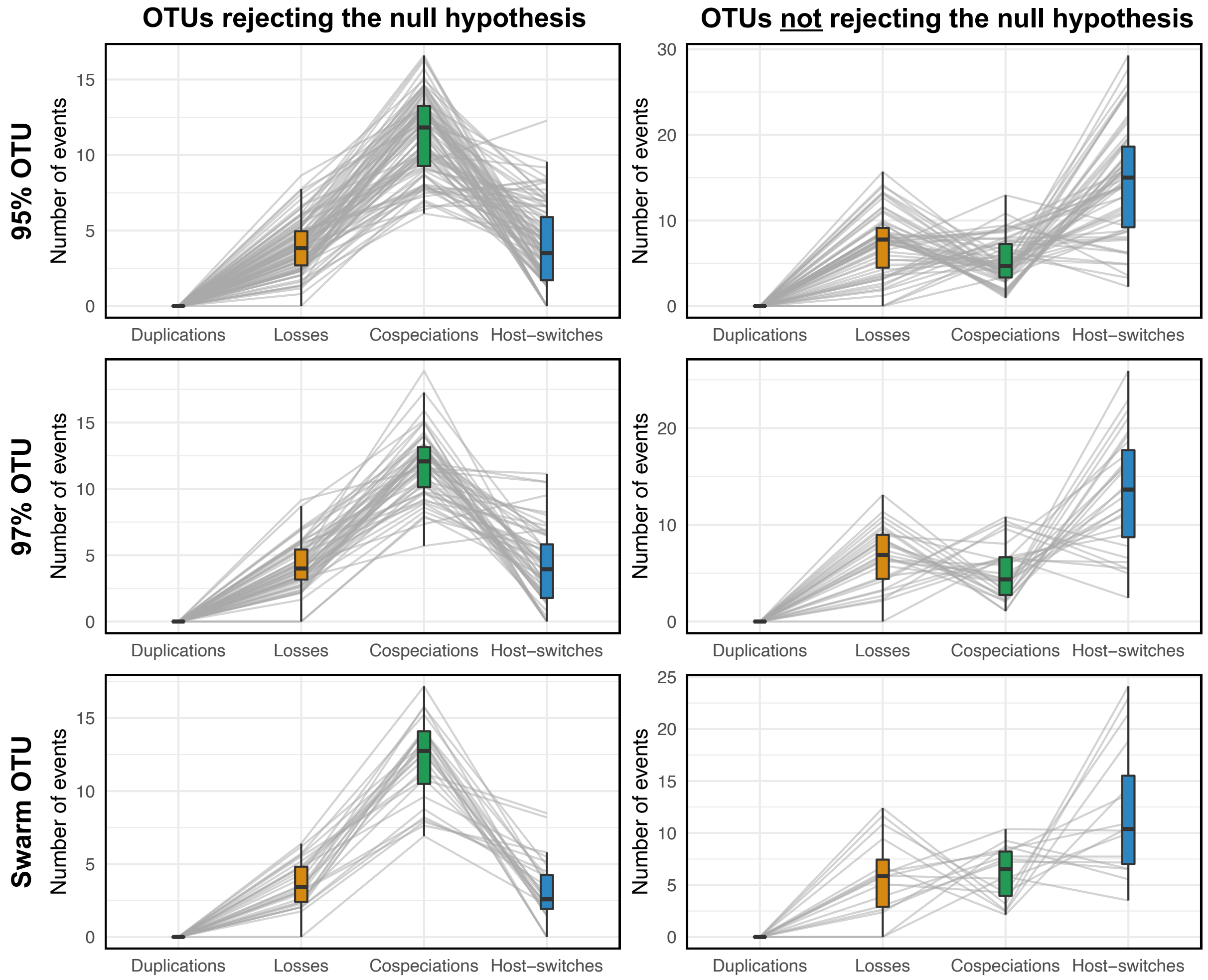

**Supplementary Figure 22: Estimated parameters for the bacterial OTUs of the primate gut microbiota using HOME**

Estimated parameters (number of host-switches or substitution rate) for various OTU criteria (either 95%, 97% sequence similarity, or Swarm OTUs), for the OTUs rejecting the null hypothesis of independent evolution (left) or not rejecting it (right).

Boxplots present the median surrounded by the first and third quartiles, and whiskers extend to the extreme values but no further than 1.5 of the inter-quartile range.

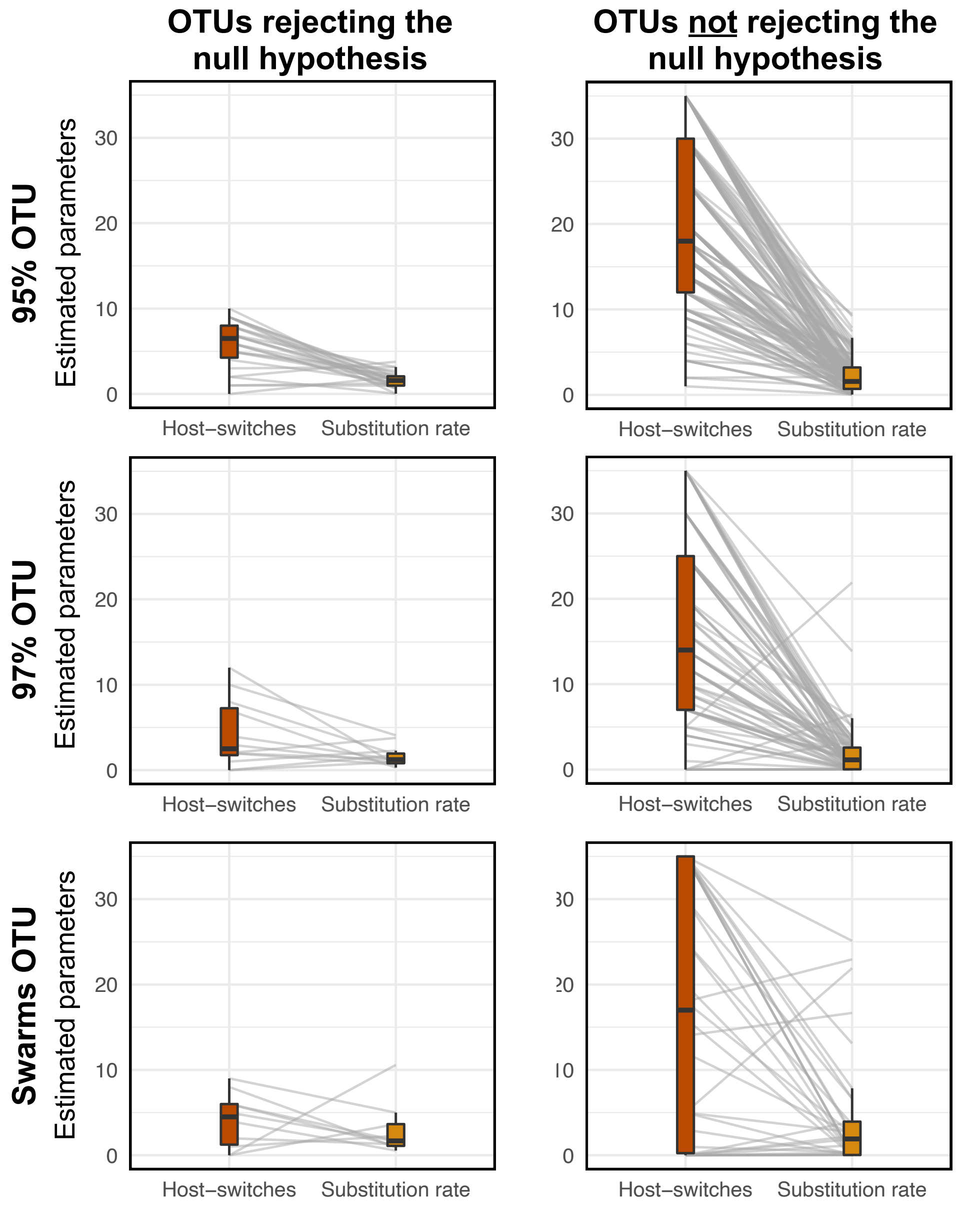

**Supplementary Figure 23: Vertical transmission in primate gut microbiota is more frequently inferred when considering several microbial strains per host species:** Numbers of OTUs from the gut microbiota of primates rejecting (in red) or not (in green) the null hypothesis of independent evolutions according to the different approaches tested: ParaFit, PACo, or ALE. In other words, OTUs colored in red represent transmitted OTUs. We did not apply HOME here as it cannot consider several microbial strains per host species. ParaFit and PACo were both evaluated using the null model 1 (top panels) and null model 2 (bottom panels).
OTUs were either clustered as 95%, 97%, or as Swarm OTUs.

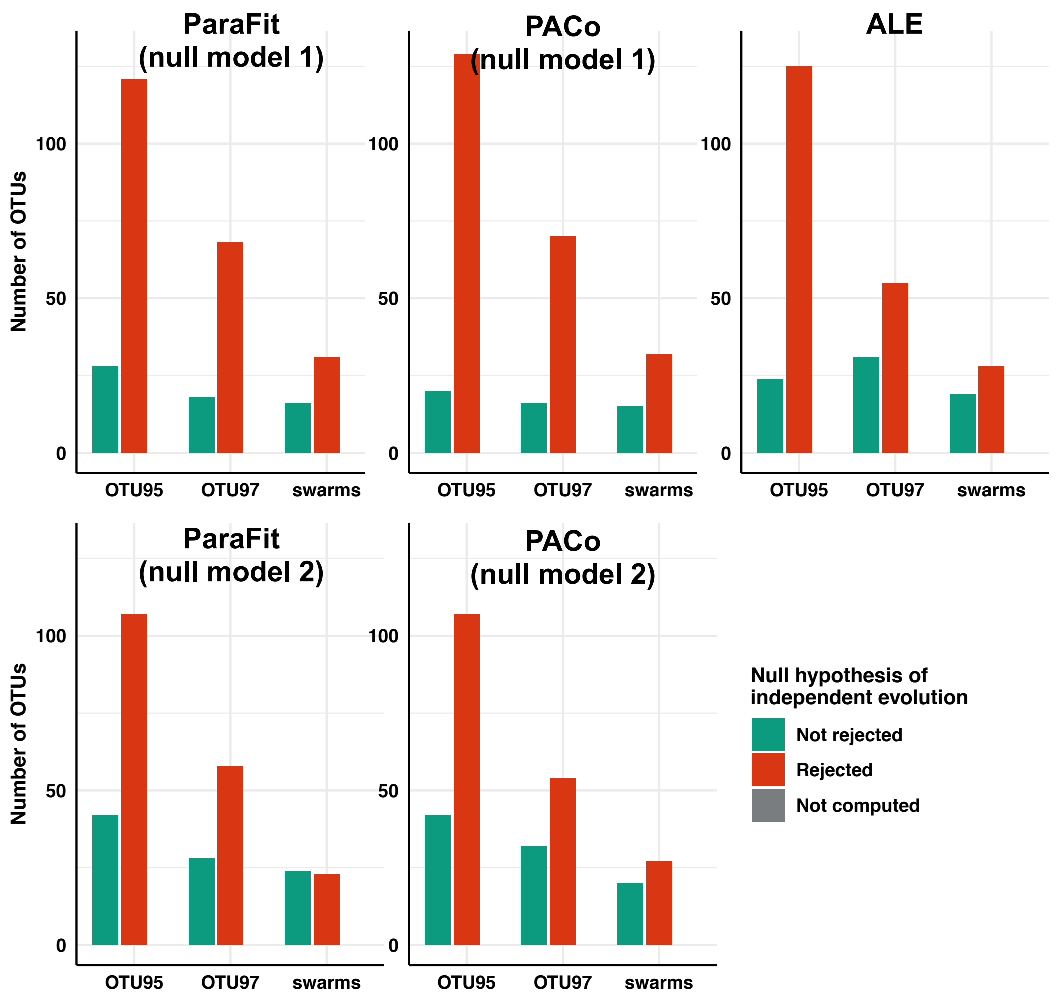
